# Characterization of two conserved cell death elicitor families from the Dothideomycete fungal pathogens *Dothistroma septosporum* and *Fulvia fulva* (syn. *Cladosporium fulvum*)

**DOI:** 10.1101/2022.06.07.495221

**Authors:** Mariana Tarallo, Rebecca L. McDougal, Zhiyuan Chen, Yan Wang, Rosie E. Bradshaw, Carl H. Mesarich

## Abstract

*Dothistroma septosporum* (Ds) and *Fulvia fulva* (Ff; previously called *Cladosporium fulvum)* are two closely related Dothideomycete fungal species that cause Dothistroma needle blight in pine and leaf mold in tomato, respectively. During host colonization, these pathogens secrete virulence factors termed effectors to promote infection. In the presence of corresponding host immune receptors, however, these effectors activate plant defenses, including a localized cell death response that halts pathogen growth. We identified two effector protein families, Ecp20 and Ecp32, which are conserved between the two pathogens. The Ecp20 family has four paralogues in both species, while the Ecp32 family has four paralogues in *D. septosporum* and five in *F. fulva.* Both families have members that are highly expressed during host infection. Members of the Ecp20 family have predicted structural similarity to proteins with a β-barrel fold, including the Alt a 1 allergen from *Alternaria alternata,* while members of the Ecp32 family have predicted structural similarity to proteins with a β-trefoil fold, such as trypsin inhibitors and lectins. Using *Agrobacterium tumefaciens-mediated* transient transformation assays, each family member was assessed for its ability to trigger cell death in leaves of the non-host species *Nicotiana benthamiana* and *N. tabacum.* Using this approach, FfEcp20-2, DsEcp20-3 and FfEcp20-3 from the Ecp20 family, and all members from the Ecp32 family, except for the Ds/FfEcp32-4 pair, triggered cell death in both species. This cell death was dependent on secretion of the effectors to the apoplast. In line with recognition by an extracellular immune receptor, cell death triggered by Ds/FfEcp20-3 and FfEcp32-3 was compromised in *N. benthamiana* silenced for *BAK1* or *SOBIR1,* which encode extracellular co-receptors involved in transducing defense response signals following apoplastic effector recognition. We then investigated whether DsEcp20-3 and DsEcp20-4 triggered cell death in the host species *Pinus radiata* by directly infiltrating purified protein into pine needles. Strikingly, as in the non-host species, DsEcp20-3 triggered cell death, while DsEcp20-4 did not. Collectively, our study describes two new candidate effector families with cell death-eliciting activity from *D. septosporum* and *F. fulva* and provides evidence that members of these families are recognized by plant immune receptors.

## 1 Introduction

Understanding how plants and fungal pathogens interact at the molecular level can lead to improvements in plant health. During infection, fungal pathogens deliver a collection of small, proteinaceous virulence factors, termed effectors, into the apoplast and/or cells of their host plants to promote the infection process (Rocafort et al., 2020; Wang et al., 2020). However, as fungal infection has a negative impact on plant fitness, there is a strong selection pressure on the host to recognize these effectors as invasion patterns and, in doing so, activate defense responses that provide protection against the pathogen (Cook et al., 2015) In line with this, in resistant hosts, particular effectors may be recognized by corresponding specific immune receptors, called resistance (R) proteins, to activate plant defense responses. Often, one of these defense responses is the hypersensitive response (HR), a localized cell death reaction that limits or halts pathogen growth (Cook et al., 2015; Win et al., 2012). As these defense responses often lead to the pathogen being unable to cause disease, such recognized effectors are typically referred to as avirulence (Avr) determinants. To circumvent recognition, and therefore the ensuing plant defense responses, the fungal invader must then modify its effector arsenal by secreting mutated or new effectors, shifting the pressure to adapt back onto the plant host, resulting in a constant “arms race” in plant-pathogen interactions (Coll et al., 2011; Jones & Dangl, 2006).

Progress in the generation of genomic, transcriptomic, and proteomic resources has enabled the rapid identification of pathogen effectors, making it possible to screen them for elicitation or suppression of plant defense responses, a process known as effectoromics. This, in turn, makes it possible to discover host defense or susceptibility genes in different plant species. A search for conserved effectors, sometimes called ‘core’ effectors, is essential as it can lead to broad spectrum resistance. The tertiary structures of some effector proteins are conserved across microbial classes because they have fundamental and important roles in infection, and for this reason are less likely to be mutated or lost during microbial evolution (Nürnberger & Brunner, 2002). Therefore, plant resistance based on the recognition of conserved effectors is more likely to be durable, as it is less likely to be overcome.

*Dothistroma septosporum* (Ds) is a hemibiotrophic pathogen that causes Dothistroma needle blight (DNB), one of the most important foliar diseases of pine trees (Drenkhan et al., 2016). Based on genomic information, *D. septosporum* is closely related to *Fulvia fulva* (Ff; syn. *Cladosporium fulvum)* (de Wit et al., 2012). *F. fulva* is a biotrophic pathogen responsible for leaf mold of tomato *(Solanum lycopersicum),* a disease that is mainly a problem in greenhouse or high tunnel environments (Thomma et al., 2005). In both the *D. septosporum*–pine and *F. fulva*–tomato pathosystems, infection begins with the germination of conidia on the leaf surface and penetration of hyphae into the host through the stomata. Once inside, both pathogens colonize the apoplastic space between mesophyll cells, where they secrete effector proteins (Kabir et al., 2015; Mesarich et al., 2018).

*F. fulva* is one of the best studied filamentous fungal pathogens. The *F. fulva*–tomato interaction is a model pathosystem in which early pioneering studies helped to establish the molecular basis of the gene-for-gene concept of plant resistance to fungal pathogens (de Wit, 2016). In *F. fulva,* small, secreted cysteine-rich proteins delivered to the host apoplast, termed extracellular proteins (Ecps) based on their identification in apoplastic wash fluid samples, have been well-studied (Mesarich et al., 2018). These include core effectors such as Ecp2-1, Ecp6 and Avr4 (de Jonge et al., 2010; Sánchez-Vallet et al., 2013; Stergiopoulos et al., 2010; van den Burg et al., 2006), homologs of which have been found in other plant-pathogenic fungi, including *D. septosporum* (Bolton et al., 2008; de Wit et al., 2012; Stergiopoulos et al., 2012). In contrast, few studies have identified and characterized effectors from *D. septosporum.* In one of these studies, DsEcp2-1, the ortholog of Ecp2-1 from *F. fulva,* was suggested to be an Avr protein responsible for eliciting defense responses in pine (Guo et al., 2020). In another of these studies, DsAvr4, the ortholog of Avr4 from *F. fulva,* was found to bind chitin, a function that is required to protect the fungal cell wall against hydrolysis by host chitinases (Mesarich et al., 2016; van den Burg et al., 2006).

In general, studies highlighting the role of effectors in molecular interactions between pathogens and gymnosperm hosts are few and far between. The close phylogenetic relationship between *D. septosporum* and *F. fulva,* however, provides a unique opportunity for comparative analysis of core effector functions across angiosperm and gymnosperm pathosystems. Recently, a set of candidate effector proteins from *D. septosporum* was transiently expressed in the non-host plants, *Nicotiana benthamiana* and *N. tabacum,* to identify those that are involved in triggering plant defense responses (Hunziker et al., 2021). Several of these proteins, all with orthologs in *F. fulva,* elicited a plant cell death response, suggesting that they are recognized by extracellular immune receptors in these non-host plants, triggering non-host resistance. Amongst these we found two proteins, Ds70057 and Ds70694, that were of particular interest, as they are orthologs of Ecps from *F. fulva* (Hunziker et al., 2021; Mesarich et al., 2018). Both proteins appear to be part of protein families – the Ecp20 family and the Ecp32 family (Hunziker et al., 2021). The Ecp20 family is made up of four members in both species, while the Ecp32 family is made up of four members in *D. septosporum* and five members in *F. fulva.*

In this study, we characterized and compared Ecp20 and Ecp32 family members from *D. septosporum* and *F. fulva* through prediction of their protein tertiary structures and by assessing their cell death-eliciting capacity in non-host angiosperm plants. We also used a novel assay involving a pine tissue culture system to determine whether two of the Ecp20 family members from *D. septosporum* trigger cell death in the gymnosperm host, *Pinus radiata.* This work provides insights into conservation of core effector function and recognition across a broad range of plant species.

## 2 Methods

### 2.1 Microorganisms and Plants

*D. septosporum* NZE10 (de Wit et al., 2012), GUA2 and COLN (Bradshaw et al., 2019), as well as *D. pini* CBS 116487 (BioSample ID SAMN02254964) and *F. fulva* 0WU (de Wit et al., 2012), were used to identify candidate effectors for characterization in this study. *Escherichia coli* DH5α (Taylor et al., 1993) and *Agrobacterium tumefaciens* GV3101 (Holsters et al., 1980) were used for gene cloning and *A. tumefaciens-mediated* transient transformation assays (ATTAs), respectively. *N. tabacum* Wisconsin 38 and *N. benthamiana* were used as model non-host plants for ATTAs, while *N. benthamiana* was used for tobacco rattle virus (TRV)-mediated gene silencing experiments. *P. radiata* clonal shoots, derived from family seed lots that are, compared to each other, relatively susceptible (S6 and S11) or tolerant (R4) to DNB, and grown from embryogenic tissue under sterile conditions on LPch agar (Hargreaves et al., 2004), were provided by Scion (New Zealand Forest Research Institute Ltd., Rotorua, New Zealand) and used in protein infiltration assays.

### 2.2 Bioinformatics

Members of the Ecp20 and Ecp32 protein families were identified from the predicted proteomes and/or genomes of *D. septosporum* NZE10 and *F. fulva* 0WU (de Wit et al., 2012). Chromosome locations of the *FfEcp20* and *FfEcp32* genes were identified using the *F. fulva* Race 5 genome assembly (Zaccaron et al., 2022). Gene family members were also identified from genome sequences of 18 other *D. septosporum* isolates (Bradshaw et al., 2019) and *D. pini* CBS 116487 (GenBank assembly accession GCA_002116355.1), using reciprocal BLASTp and tBLASTn searches in conjunction with an E-value cut-off of <10^-5^. BLASTp was also employed to determine the level of conservation for each family member across fungal species in the Joint Genome Institute (JGI) MycoCosm database, using an E-value threshold of 10^-5^. EffectorP v3.0 (Sperschneider & Dodds, 2021) was used to predict whether members of the Ecp20 and Ecp32 families are effectors. All *D. septosporum* NZE10, *D. pini* CBS 116487 and *F. fulva* 0WU nucleotide and amino acid sequences for Ecp20 and Ecp32 family members are shown in Supplementary Table S1.

Protein tertiary structure predictions were performed using AlphaFold2 (Jumper et al., 2021; Mirdita et al., 2022), and the Dali server (Holm, 2020) was used to identify proteins with structural similarity in the Research Collaboratory for Structural Bioinformatics Protein Data Bank (RCSB PDB). Here, a Dali Z-score of 2 or greater was used to infer structural similarity. Protein tertiary structures were visualized and rendered in PyMol v2.5, with alignments carried out using the CEalign tool (DeLano, 2002).

Protein sequence alignments were performed using Clustal Ω (Sievers et al., 2011) in Jalview (www.jalview.org). Phylogenetic trees were constructed with the neighbor-joining method using Geneious Software v9.1.8 (Kearse et al., 2012).

### 2.3 ATTA Expression Vectors

Candidate effector genes of *D. septosporum* NZE10, GUA2 and COLN, *D. pini* CBS 116487 and *F. fulva* 0WU were cloned in the pICH86988 ATTA expression vector, which contains the nucleotide sequence encoding the *N. tabacum* PR1α signal peptide for secretion into the plant apoplast, followed by a 3×FLAG for detection via Western blotting (Weber et al. (2011), according to the method described by Guo et al. (2020) in cases where the genes had no predicted introns, or synthesized directly into pICH86988 by Twist Bioscience (San Francisco, CA, USA) in cases where the genes had predicted introns. For the genes that were cloned into pICH86988, the mature coding sequence was first amplified by polymerase chain reaction (PCR) from genomic DNA extracted from the fungus, with genomic DNA extraction performed as described by Doyle and Doyle (1987). PCRs were performed with Phusion High-Fidelity DNA polymerase (New England Biolabs, Beverly, MA, USA), and PCR products were extracted from agarose gels using an E.Z.N.A. Gel Extraction Kit (Omega Bio-tek, Norcross, GA, USA). PCR primers were designed with Primer3 (Untergasser et al., 2012) in Geneious v9.1.8 (Kearse et al., 2012). These primers, which were synthesized by Integrated DNA Technologies (IDT; Supplementary Table S2), had 5’ extensions containing a BsaI recognition site and 4-base overhang specific for Golden Gate modular cloning (Engler et al., 2009) in between the CaMV 35S promoter and octopine synthase terminator of pICH86988. Versions of the candidate effector genes were also ligated into pICH86988 without the nucleotide sequence encoding the PR1α signal peptide to prevent secretion of the proteins they encode into the apoplast. Constructs were then transformed into *E. coli,* plasmids extracted using an E.Z.N.A.® Plasmid DNA Mini Kit I (Omega Bio-tek), and correct assemblies confirmed by sequencing. ATTA expression vectors were transformed into *A. tumefaciens* by electroporation as described previously (Guo et al., 2020).

### 2.4 ATTAs

To heterologously express candidate effector genes in *N. benthamiana* and *N. tabacum* using ATTAs, single colonies of *A. tumefaciens* carrying pICH86988 constructs of interest were first inoculated into selective lysogeny broth (LB) containing 50 μg/mL kanamycin, 10 μg/mL rifampicin and 30 μg/mL gentamycin, then incubated overnight at 28°C. Cells were subsequently collected by centrifugation at 2,500 *g* for 5 min and resuspended in 1 mL of infiltration buffer (10 mM MgCl_2_:6H_2_O, 10 mM MES-KOH (Sigma-Aldrich, St. Louis, MO, USA), 100 μM acetosyringone (Sigma-Aldrich)) to an OD_600_ of 0.5. Following resuspension, the abaxial surface of *N. benthamiana* and *N. tabacum* leaves was infiltrated with the appropriate *A. tumefaciens* suspension using 1 mL needleless syringes. Here, the extracellular elicitin INF1 from *Phytophthora infestans,* which is recognized by the LRK1 immune receptor in *Nicotiana* species (Kamoun et al., 1997; Kanzaki et al., 2008), was used as a positive cell death control, while the empty pICH86988 vector was used as a negative no cell death control. At least 12–24 infiltration zones were tested for each candidate effector gene or control (consisting of two leaves from each of two plants ×3–6 repeat experiments). Plants were monitored for up to 7 days, and photographs taken using a Nikon D7000 camera. Infiltration zones were assessed on whether the protein tested elicited a strong or weak cell death response, or no cell death. A strong cell death response was scored when it was indistinguishable from the response elicited by INF1 and around the entire infiltration zone, while a weak response was considered when the protein elicited a weaker response around approximately 50% of the infiltration zone when compared with INF1. A no-cell death response was indicated when the infiltration zone had the same response as the negative empty pICH86988 vector control.

### 2.5 Western Blotting

To confirm that candidate effector proteins had been produced in *N. benthamiana* and *N. tabacum* leaves using ATTAs, and to determine whether these proteins were of the predicted size, Western blotting was conducted with total protein as described previously by Guo et al. (2020). For protein separation by SDS-PAGE, 12% separating and 5% stacking gels were used, with proteins transferred to PVDF membranes. Anti-FLAG® M2 primary antibody produced in mouse (Sigma-Aldrich), together with anti-mouse secondary antibody produced in chicken (Santa Cruz Biotechnology, Dallas, TX, USA) and SuperSignal® West Dura Extended Duration substrate (Thermo Fisher Scientific, Waltham, MA, USA), were used for protein detection. Proteins were visualized using an Azure Biosystems c600 Bioanalytical Imaging system (Azure Biosystems, Dublin, CA, USA).

### 2.6 Protein Expression and Purification

Genes encoding candidate effector proteins were amplified by PCR as above from the previously generated ATTA expression vectors, without their native signal peptide sequences. Here, the 5’ PCR primers incorporated a FLAG-tag for detection by Western blotting. PCR primers are shown in Supplementary Table S2. PCR amplicons were gel-extracted as above and ligated, using DNA ligase (New England Biolabs), into the SmaI/EcoRI restriction sites of the expression vector pPic9-His_6_ (Invitrogen, Carlsberg, CA, USA), which encodes an α-factor signal peptide for protein secretion and a hexa-histidine (6×His) tag for protein purification. The ligated products were then transformed into *E. coli.* The expression constructs were sequenced and contained the predicted mature sequence of the candidate effectors fused to a 6×His tag at their N-terminus. The expression cassette was linearized with SacI or SalI (New England Biolabs) restriction enzymes and transformed into *P. pastoris* GS115, according to Kombrink (2012).

Protein expression in *P. pastoris* was performed according to Weidner et al. (2010). Briefly, protein expression was induced in 200 mL of BMMY (Buffered Methanol-complex Medium), through the addition of methanol to a concentration of 0.5% (v/v) every 24 h, for 72 h. The cells were then removed by centrifugation at 4,500 *g* for 30 min and the supernatant filter-sterilized by passage through a 0.22 μm membrane (ReliaPrep, Ahlstrom, Helsinki, Finland). Finally, the pH of the sterilized culture filtrate was adjusted to 8 with NaOH.

The secreted candidate effector protein (fused with an N-terminal 6×His tag) was purified from culture filtrate using immobilized metal ion affinity chromatography (IMAC), in conjunction with a Ni Sepharose™ 6 Fast Flow (GE Healthcare, Chicago, IL, USA) column, according to the manufacturer’s protocol. Before loading, the column (Glass Econo-Column – BioRad, Hercules, CA, USA) was packed with 5 mL of resin and equilibrated by washing with binding buffer (20 mM sodium phosphate, 0.5 M NaCl, pH 7.4). The culture filtrate was added to the column at a flow rate of 1 mL/min, and protein eluted with an elution buffer (20 mM sodium phosphate, 0.5 M NaCl and 500 mM imidazole, pH 7.4). The elution fractions were mixed, and elution buffer was added to a final volume of 50 mL to obtain sufficient volume for vacuum-infiltration of pine shoots. Western blotting was performed as above to determine the presence of each candidate effector protein in *P. pastoris* culture filtrate and after purification. Protein concentrations in culture filtrates and purified solutions were determined by using a Pierce™ Coomassie (Bradford) Protein Assay Kit (Thermo Fisher Scientific), as per the manufacturer’s protocol.

### 2.7 Pine Infiltration with Purified Proteins

Proteins were infiltrated into tissue-cultured pine shoots at a concentration of approx. 20 μg/mL using the method developed by Hunziker et al. (2021). Briefly, clonal rootless pine shoots were maintained in LPch agar (Hargreaves et al., 2004) in glass jars, with each jar containing 6–8 shoots. For this experiment, clonal shoots originating from two relatively *Dothistroma*-susceptible families (S6 and S11) and one relatively *Dothistroma-tolerant* family (R4) (based on comparative field data) were used. The shoots were completely submerged in approx. 50 mL of each purified candidate protein or control solution (positive control: DsEcp32-3; negative control: elution buffer). Samples were exposed to vacuum in a glass chamber for 5 min to facilitate protein uptake. Shoots were then briefly rinsed in sterile MilliQ (MQ) water and subsequently placed back into the LPch agar in the glass jar. The experiment was conducted at 22°C in a room with natural light. Photos were taken 7 days after infiltration (dai) using a Nikon D7000 camera.

### 2.8 Tobacco Rattle Virus (TRV)-mediated Virus-induced Gene Silencing (VIGS)

Fragments used to generate virus-induced gene silencing (VIGS) constructs were amplified by PCR from cDNA or genomic DNA of *N. benthamiana* using PrimeSTAR GXL DNA Polymerase (Takara Bio Inc.) with the primers listed in Supplementary Table S2. The TRV VIGS system is composed of pTRV-RNA1, which encodes replication related proteins, and pTRV-RNA2, which harbors the viral coat protein, and the candidate fragment sequences that target silencing to the corresponding genes (Liu et al., 2002; MacFarlane, 1999). The purified fragments were cloned into the pTRV-RNA2 vector (Dong et al., 2007) using the ClonExpress II One Step Cloning Kit (Vazyme Biotech Co., Ltd, Nanjing, China). The resulting vectors were verified by sequencing and individually transformed into *A. tumefaciens* strain. DsEcp20-3 was screened against around 100 TRV VIGS constructs which targeted different *N. benthamiana* LRR-RLP and LRR-RLK-encoding genes. Information on this library of silencing constructs can be found in Supplementary Data 1 from Wang et al. (2018). Primers for *T26, T26-2,* TRV:*BAK1* (Heese et al., 2007) and TRV:*SOBIR1* (Liebrand et al., 2013) are shown in Supplementary Table S2.

*A. tumefaciens* strains carrying *N. benthamiana* gene silencing vectors (Wang et al., 2018) were grown overnight in LB medium with the appropriate antibiotics. *A. tumefaciens* cells were pelleted as mentioned above, resuspended, and incubated in infiltration medium (10□mM 2-[N-morpholino] ethanesulfonic acid pH 5.6, 10□ mM MgCl_2_, 200□μM acetosyringone) for 3–4□h before agroinfiltration. *A. tumefaciens* cultures expressing pTRV-RNA2 constructs were mixed in a 1:1 ratio with *A. tumefaciens* culture expressing pTRV1 to a final OD_600_ of 1.0 before infiltration into cotyledons of four-leaf-stage *N. benthamiana* seedlings. *P. sojae* NPP1 and XEG1 (Fellbrich et al., 2002; Ma et al., 2015), and *P. infestans* INF1 were used as cell death-eliciting controls for VIGS. XEG1- and INF1-triggered cell death was known to be compromised in *N. benthamiana* leaves treated with TRV:*BAK1* or TRV:*SOBIR1*, but not with TRV:*GFP*, which was used as a negative control. (Wang et al., 2018).

### 2.9 RNA Isolation and Quantitative Reverse Transcription PCR

To determine the expression levels of *BAK1, SOBIR1* and *T26* in the TRV-mediated VIGS experiment, *N. benthamiana* leaves of similar size were selected and ground to a powder in liquid nitrogen. Total RNA was isolated from 100 ? mg of ground material using the E.Z.N.A.® Total RNA Kit I (Omega Bio-tek) and used as a template for first strand cDNA synthesis with PrimeScript Reverse Transcriptase (Takara Bio Inc., Kusatsu, Shiga, Japan). Quantitative reverse transcription PCR (qRT-PCR) was performed on an ABI 7500 Fast Real-Time PCR system (Applied Biosystems Inc., Foster City, CA, USA) using PrimeScript RT Master Mix (Takara Bio Inc.) with translation elongation factor 1 alpha *(EF-1α)* as an endogenous control and the primers listed in Supplementary Table S2. Data were analyzed using the 2^-ΔΔCT^ method (Livak & Schmittgen, 2001).

## 3 Results

### 3.1 Two *D. septosporum* Elicitors are Effector Candidates that Form Part of Protein Families with Orthologs in *F. fulva*

In our previous work, two small, secreted candidate effector proteins from *D. septosporum* NZE10, Ds70694 and Ds70057, triggered cell death in the non-host species *N. tabacum* and *N. benthamiana* (Hunziker et al., 2021). Both proteins are homologs of Ecps from *F. fulva* and are part of two different protein families, Ecp20 and Ecp32, respectively (Mesarich et al., 2018). More specifically, Ds70694 is an ortholog of FfEcp20-3 from *F. fulva* and was therefore named DsEcp20-3, whilst Ds70057 is an ortholog of FfEcp32-3 and was therefore named DsEcp32-3.

The Ecp20 family has four paralogues in both species. Based on sequence similarities between them, DsEcp20-3 and FfEcp20-3, together with the other orthologous pairs Ds/FfEcp20-1, Ds/FfEcp20-2 and Ds/FfEcp20-4, were grouped together (Figure 1A, Supplementary Figure S1A). Interspecies pairwise amino acid identities between all family members from *D. septosporum* and *F. fulva* ranged from 10.6% to 85.5%, while intraspecies pairwise identities between all family members ranged from 14.2% to 40.9% in *D. septosporum* and 11.8% to 44.2% in *F. fulva* (Supplementary Table S3). The *Ds/FfEcp20-1* genes contain one exon, with the intron located at the same position, while genes *Ds/FfEcp20-2, −3* and *−4* do not have any introns (Supplementary Figure 2A). All genes encode a protein with four cysteine residues, although the location of these varies slightly from protein pair to protein pair (Supplementary Figures S1A and S2A).

**Figure 1.**
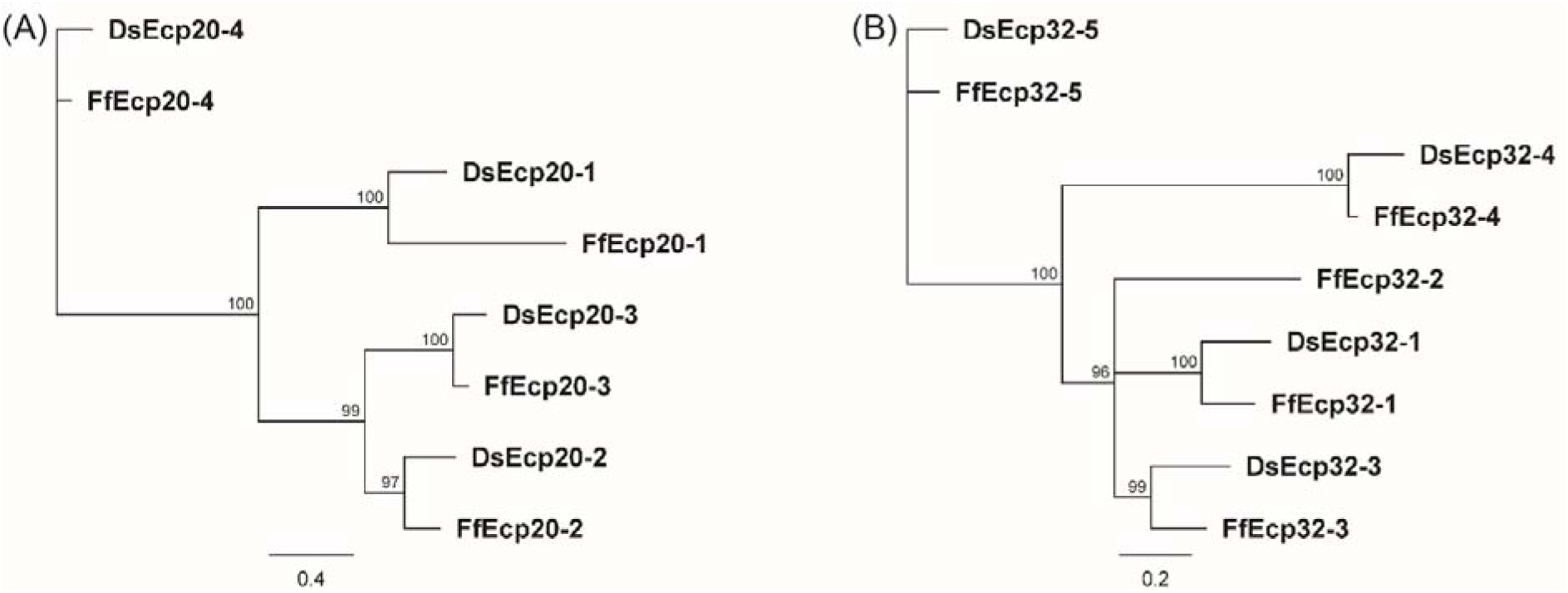
Protein phylogenies of the *Dothistroma septosporum* and *Fulvia fulva* Ecp20 (A) and Ecp32 (B) families. The trees were constructed with the neighbor-joining method using Geneious Software v9.1.8. Bootstrap values are shown at the nodes as percentages. The scale bar represents 0.4 (A) and 0.2 (B) substitutions per site. Ds: *D. septosporum;* Ff: *F. fulva.*

Based on data from a previously published transcriptome study (Bradshaw et al., 2016), all genes of the *D. septosporum Ecp20* family are expressed during pine infection, with *DsEcp20-3* and *DsEcp20*-4 being the two most highly expressed (Supplementary Table S4). *DsEcp20-3* showed strong up-regulation in the Mid and Late *in planta* stages that mark the start and end of necrotrophic infection, when compared with the Early (biotrophic) stage and *in vitro,* while *DsEcp20-4* is strongly up-regulated in the Late *in planta* stage. For the corresponding genes in *F. fulva* during tomato infection, as based on data from a previously published transcriptome study (Mesarich et al., 2018), *FfEcp20-2* and *FfEcp20-3* were highly expressed throughout the timepoints analyzed, while *FfEcp20-4* was highly expressed at the beginning of infection and downregulated at later time points, when compared with *in vitro* (Supplementary Table S4). It should be noted that *FfEcp20-1* had no gene model and therefore had no FPKM value assigned to it in the previous study (Mesarich et al., 2018). Thus, no expression information was available for this gene.

The Ds/FfEcp32 family was analyzed in the same way as the Ds/FfEcp20 family above. The Ecp32 family has five paralogues in *F. fulva,* named FfEcp32-1 to −5, and four in *D. septosporum,* named DsEcp32-1 and DsEcp32-3 to −5 (there is no ortholog of FfEcp32-2 in *D. septosporum)* (Figure 1B, Supplementary Figure S1B). Interspecies pairwise amino acid identity between all family members from *D. septosporum* and *F. fulva* ranged from 10.6% to 82%, while intraspecies pairwise identity between members ranged from 22.1% to 42.1% in *D. septosporum* and 21.6% to 49% in *C. fulvum* (Supplementary Table S3). Regarding the exon-intron structure, *Ds/FfEcp32-1, FfEcp32-2* and *Ds/FfEcp32-3* do not have any introns. In contrast, the *Ds/FfEcp32-4* gene pair has one intron at the same position, while the *Ds/FfEcp32-5* gene pair has two introns at the same positions (Supplementary Figure S2B). In terms of cysteine residues, the Ds/FfEcp32 proteins have anywhere from none (Ds/FfEcp32-5) to five (Ds/FfEcp32-4), although in the case of DsEcp32-3, an additional cysteine is present compared to the Ff ortholog (Supplementary Figures S1B and S2B).

All *DsEcp32* genes are also expressed during pine infection. However, only *DsEcp32-3* and *DsEcp32-5* are highly expressed *in planta,* especially at the Mid and Late necrotrophic stages when compared with *in vitro. DsEcp32-1* and *DsEcp32-4* are only expressed at a low level *in planta.* During *F. fulva* infection of tomato, *FfEcp32-1, FfEcp32-2* and *FfEcp32-3* are highly expressed at all timepoints. *FfEcp32-4* was not expressed under any condition, while *FfEcp32-5* was only expressed at 12 dpi, and was downregulated compared with *in vitro* (Supplementary Table S4).

Genomes from the two species are mesosyntenic (de Wit et al., 2012; Zaccaron et al., 2022) and, in keeping with this, we found conservation of chromosome location between the two species for each of the *Ds/FfEcp20* and *Ds/FfEcp32* gene pairs (Supplementary Figure S3). Taken together, sequence similarity, intron position and chromosome localization suggest a common evolutionary origin for the *Ecp20* and *Ecp32* genes from both species.

### 3.2 Predicted Tertiary Structures of Ecp20 and Ecp32 Family Members Suggest Possible Roles in Virulence

We predicted the tertiary structures of all Ecp20 family members from *D. septosporum* NZE10 and *F. fulva* 0WU using AlphaFold2 (Figure 2; Supplementary Figure S4). All proteins from the Ecp20 family are predicted to be structurally similar to the characterized β-barrel fold of Alt a 1 from *Alternaria alternata* (RCSB protein data bank [PBD] ID: 3V0R) (Chruszcz et al., 2012) (Supplementary Table S4 and S5). Figure 2 shows the predicted structures for DsEcp20-3 and DsEcp20-4, as well as the alignments between DsEcp20-3 and the characterized structure of Alt a 1 from *A. alternata* and between the predicted structures of DsEcp20-3 and DsEcp20-4. The predicted structures for all the DsEcp20 and FfEcp20 proteins are shown in Supplementary Figure S4 along with characterized crystal structures of similar proteins from other fungal species.

**Figure 2.**
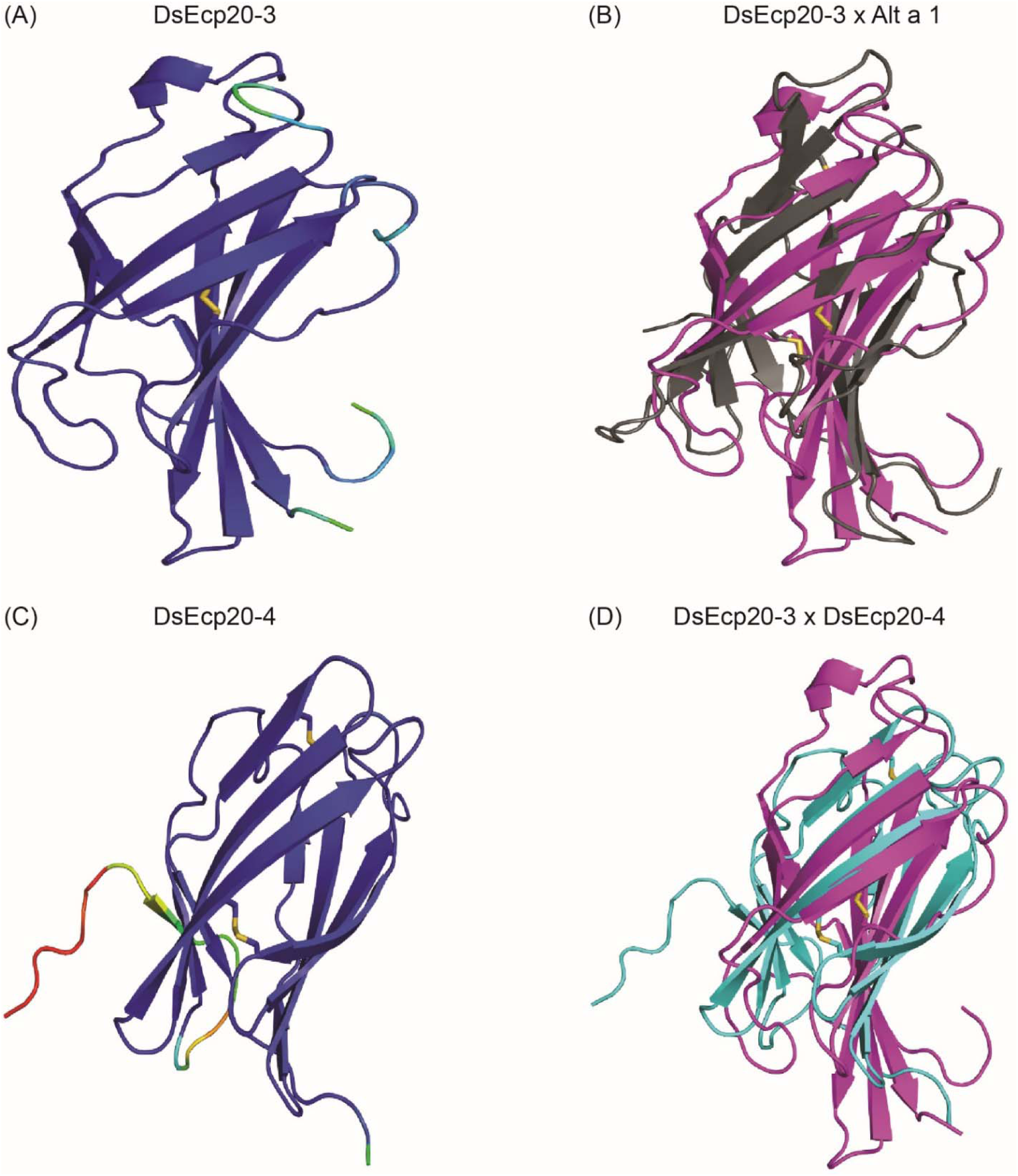
Predicted protein tertiary structures of Ecp20 family members from *Dothistroma septosporum.* (A) Predicted structure of DsEcp20-3. (B) Alignment of the predicted DsEcp20-3 structure (pink) with the characterized structure of Alt a 1 (black) from *Alternaria alternata* (RCSB protein data bank [PBD] ID: 3V0R) (Chruszcz et al., 2012). (C) Predicted structure of DsEcp20-4. (D) Alignment of predicted DsEcp20-3 (pink) and DsEcp20-4 (blue) structures. Disulphide bonds are shown as yellow sticks. Protein structures were predicted with AlphaFold2, rendered in PyMol v2.5 and aligned using the CEalign tool (DeLano, 2002; Jumper et al., 2021; Mirdita et al., 2022). Structures on the left were colored in PyMol according to their AlphaFold2 pLDDT score: dark blue for regions predicted with high confidence, light blue and green for regions of low confidence, and red for very low confidence regions.

The tertiary structure of Ecp32 proteins was also predicted using AlphaFold2 (Figure 3, Supplementary Figure S5). All Ecp32 family members had significant hits with trypsin inhibitors and lectins; these are proteins that share a β-trefoil fold and are known virulence factors in plant-pathogenic fungi (Renko et al., 2012; Sabotic et al., 2019; Varrot et al., 2013) (Supplementary Table S5). Figure 3 shows the predicted structures for DsEcp32-3 and DsEcp32-4, as well as the alignments between DsEcp32-3 and the characterized structure of a trypsin inhibitor from the plant species *Enterolobium contortisiliquum* (RCSB PBD ID: 4J2K) (Zhou et al., 2013), which is the top hit in the PDB for most Ds/FfEcp32 proteins. The alignment between the predicted structures of DsEcp32-3 and DsEcp32-4 is also shown in Figure 3, while all the predicted Ds/FfEcp32 structures can be found in Supplementary Figure S5 along with characterized structures of similar proteins.

**Figure 3.**
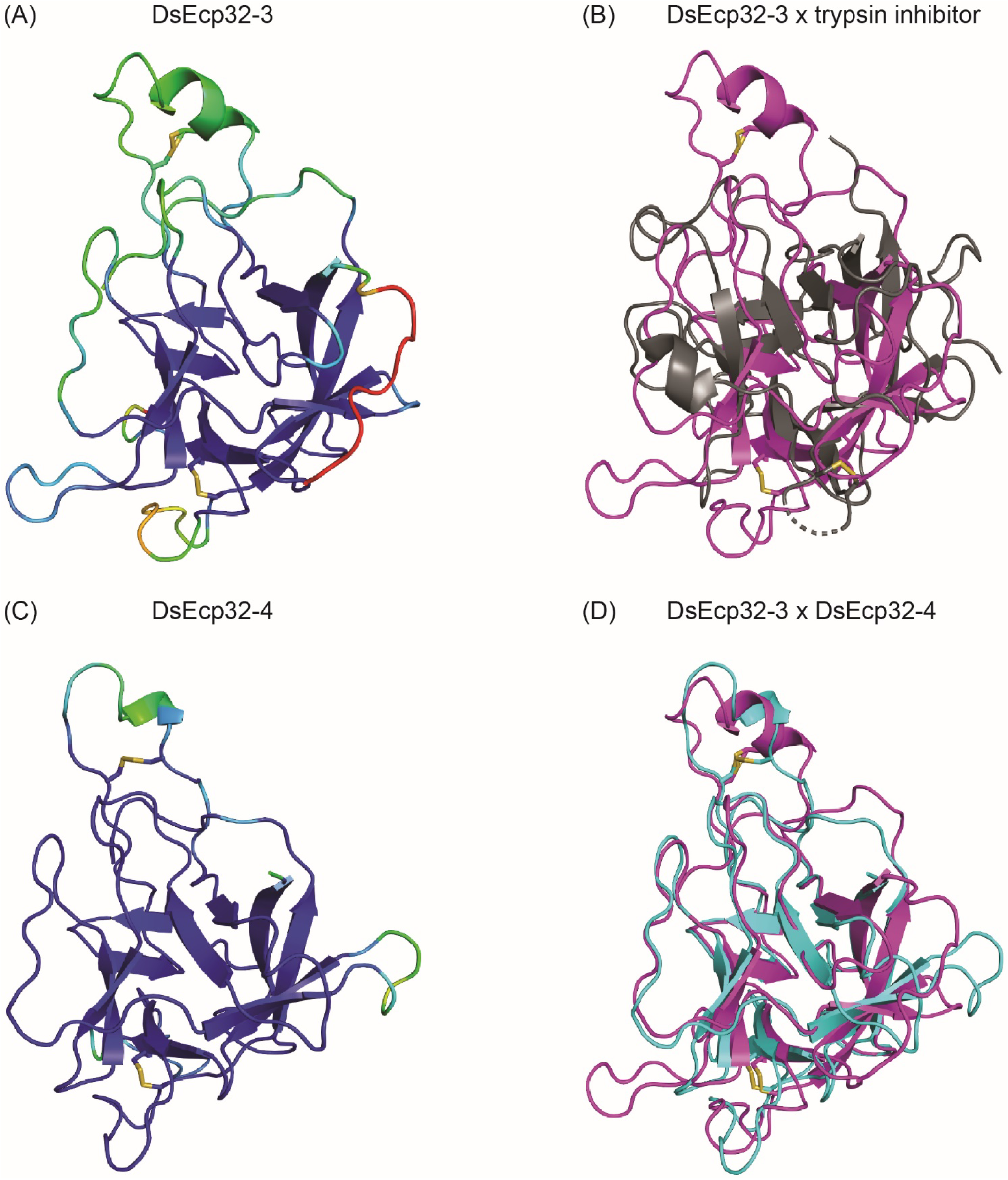
Predicted protein tertiary structure of Ecp32 family members from *Dothistroma septosporum.* (A) Predicted structure of DsEcp32-3. (B) Alignment of the predicted DsEcp32-3 (pink) structure with the characterized structure of trypsin inhibitor (black) from *Enterolobium contortisiliquum* (RCSB protein data bank [PBD] ID: 4J2K) (Zhou et al., 2013). (C) Predicted structure of DsEcp32-4. (D) Alignment of predicted DsEcp32-3 (pink) and DsEcp32-4 (blue) structures. Disulphide bonds are shown as yellow sticks. Protein structures were predicted with AlphaFold2, rendered in PyMol v2.5 and aligned using the CEalign tool (DeLano, 2002; Jumper et al., 2021; Mirdita et al., 2022). Structures on the left were colored in PyMol according to their AlphaFold2 pLDDT score: dark blue for regions predicted with high confidence, light blue and green for regions of low confidence, and red for very low confidence regions.

### 3.3 Ecp20 and Ecp32 Family Members are Conserved Across Other Fungal Species and *D. septosporum* isolates

The Ecp20 family is present in fungal species from other classes. DsEcp20-1 had 27 significant BLASTp hits in the JGI MycoCosm protein database. Two of these were from other Dothideomycete species (including the ortholog in *F. fulva),* while the rest were with from species in the Eurotiomycetes class (Supplementary Table S6). DsEcp20-2 and DsEcp20-3, the most closely related to each other among the DsEcp20 family members (Figure 1A), both had the same 28 hits in JGI. All of these matches were with other Dothideomycete species and, of these, 24 were hits to proteins from other plant pathogens (Supplementary Table S6). DsEcp20-4 had 1,006 hits in the JGI MycoCosm protein database, of which 241 were proteins from plant pathogens (Supplementary Table S6).

In general, the Ecp32 family is more widely conserved among other fungal species compared to the Ecp20 family. Individual searches in the JGI MycoCosm protein database for each of the DsEcp32 family members yielded approximately 1,000 hits for each family member (Supplementary Table S6). Those hits included proteins from fungal species with different lifestyles, ranging from saprophytes, to biotrophs, hemibiotrophs and necrotrophs. For example, DsEcp32-3 had 158 hits to proteins from fungal plant pathogens in JGI. Of these, 42 were with other members from the Mycosphaerellaceae family. These included *Cercospora zeae-maydis* with four paralogs, *Pseudocercospora (Mycosphaerella) eumusae* with six, *Pseudocercospora fijiensis* with five and *Zymoseptoria tritici* with three paralogs (Supplementary Table S6).

*Dothistroma pini* is also a causal agent of DNB (Barnes et al., 2004). Thus, we searched for orthologs of DsEcp20/32 encoded in the *D. pini* genome. All proteins from the *D. septosporum* Ecp20 and Ecp32 families had orthologs in the genome of *D. pini,* except Ecp32-5, for which the gene encoding this protein appears to be a pseudogene. The protein sequences were very similar between family members from the two *Dothistroma* species, with full-length pairwise amino acid identity varying from 81.1% to 93.8% (Supplementary Tables S1 and S3).

We next addressed the question of whether the *D. septosporum Ecp20* and *Ecp32* family genes are conserved in other isolates of this species and whether they are identical in sequence to NZE10. Genome sequences were available from 18 *D. septosporum* isolates collected from different geographic locations (Bradshaw et al., 2019). All members of the *D. septosporum Ecp20* and *Ecp32* families were present in the 18 other genomes. In each of these genomes, there was at least one single nucleotide polymorphism (SNP) leading to an amino acid difference in at least one family member for both Ecp20 and Ecp32 (Supplementary Table S7).

A total of 59 different SNPs were identified among the *Ecp20* family members across the 18 *D. septosporum* isolates, compared to NZE10 (Figure 4). *DsEcp20-1*, *DsEcp20-2* and *DsEcp20-4* had approximately the same number of SNPs: 11, 13 and 15, respectively, while *DsEcp20-3* had a total of 20. For *DsEcp20-1, DsEcp20-3* and *DsEcp20-4,* however, most SNPs (61%, 60% and 60%, respectively) were synonymous. In the case of *DsEcp20-2*, however, only 30% were synonymous and a higher number of SNPs leading to an amino acid difference (9 out of 13; 70%) were observed (Figure 4A). The *Ecp32* gene family had a higher number of DNA modifications across the 18 *D. septosporum* genomes analyzed, with a total of 117 SNPs and 3 insertions. The two most polymorphic genes were *DsEcp32-3* and *DsEcp32-5*. Interestingly, while *DsEcp32-3* had 31 sequence modifications, only five of these led to an amino acid difference (16%). All other family members had 9 or 10 SNPs leading to an amino acid difference (Figure 4B).

**Figure 4.**
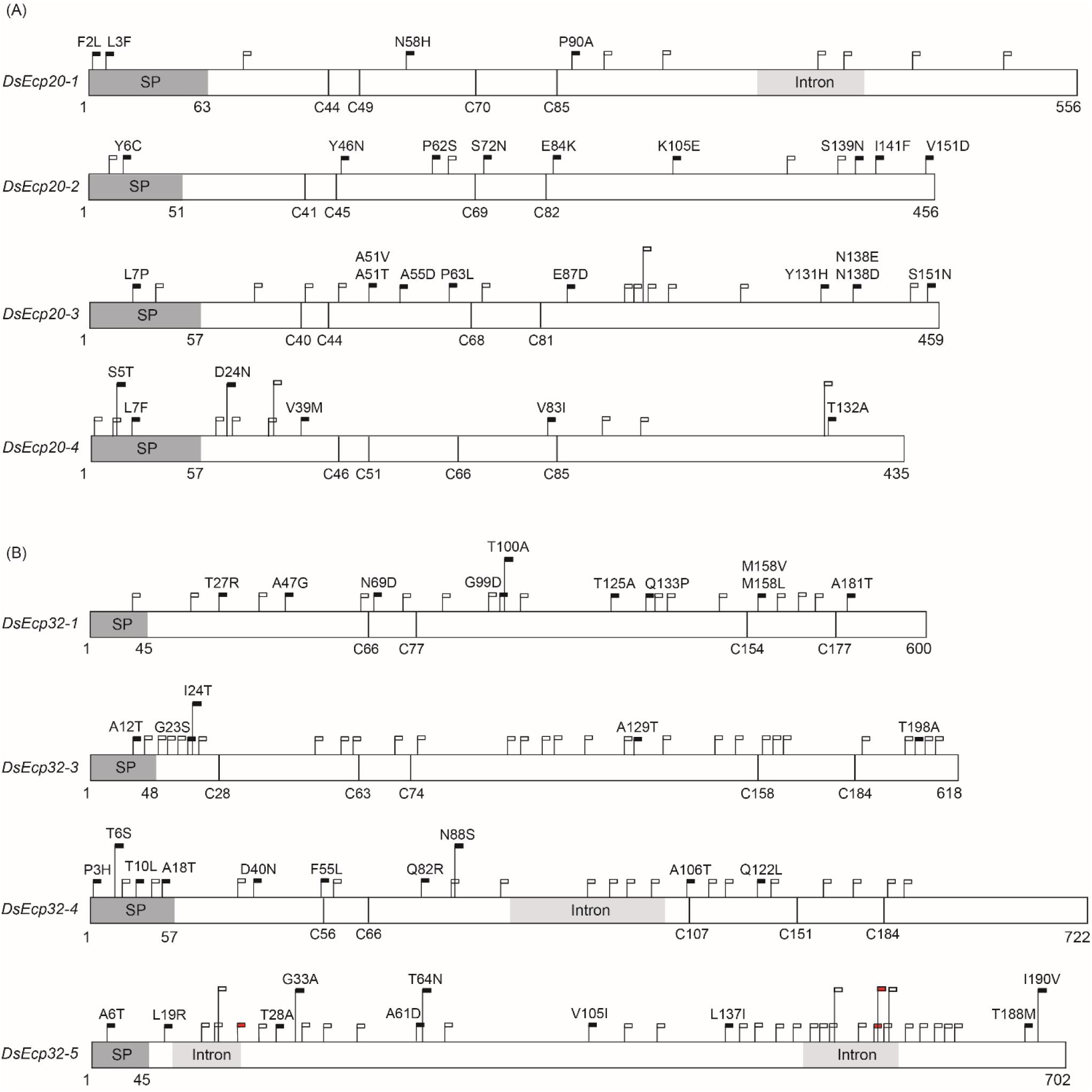
Schematic diagram showing sequence variation in *Dothistroma septosporum* NZE10 *Ecp20* (A) and *Ecp32* (B) family members across 18 other *D. septosporum* isolates. Dark grey boxes indicate signal peptide positions and light grey boxes indicate intron positions. Black flags show the position of nucleotide variations that led to non-synonymous changes, with the specific amino acid differences indicated above the flags. White flags show the position of synonymous nucleotide variations. Red flags show the position of nucleotide insertions. Vertical black lines show the positions of cysteine residues. Nucleotide positions are shown under each gene. SP: signal peptide.

The Guatemalan (GUA1 and GUA2) and Colombian (COLN and COLS) isolates showed the greatest amino acid sequence diversity, when compared to strain NZE10, for both the Ecp20 and Ecp32 family members. For DsEcp20-1, the most diverse isolates were COLN and COLS, with identical sequences, while COLN had the most sequence variation in DsEcp32-1. For DsEcp20-2 and −4, and Ecp32-3, −4, and −5, both Guatemalan isolates were the most diverse and, for DsEcp20-3, GUA2 had greater amino acid diversity (Supplementary Table S7).

### 3.4 Other Members of the Ecp20 and Ecp32 Families from *D. septosporum* and *F. fulva* Trigger Cell Death Responses in Non-host Plants

Following on from a previous study in which DsEcp20-3 and DsEcp32-3 were shown to elicit cell death in the non-host plants *N. benthamiana* and *N. tabacum* using ATTAs (Hunziker et al., 2021), we set out to determine whether other family members of both *D. septosporum* and *F. fulva* could elicit the same response. To achieve this, all Ecp20 and Ecp32 family members were transiently expressed in *N. benthamiana* and *N. tabacum* using ATTAs, and the percentages of plant responses in three different categories recorded: strong cell death, weak cell death and no cell death (Figure 5; Supplementary Figure S6). Here, orthologous pairs from both pathogens were tested on the same leaf, so that differences between them could be better assessed.

**Figure 5.**
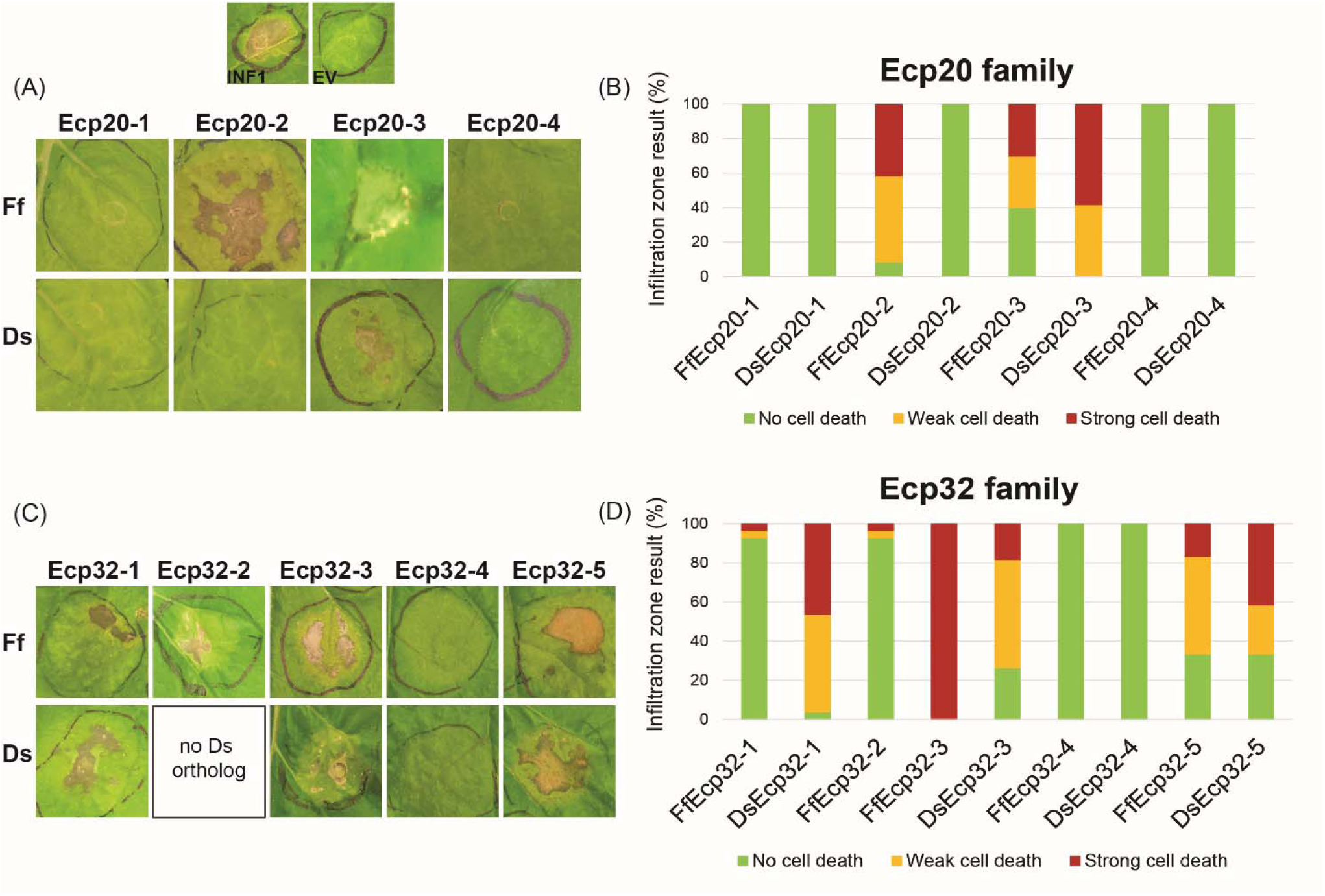
Ecp20 and Ecp32 proteins from *Dothistroma septosporum* NZE10 and *Fulvia fulva* 0WU trigger cell death responses in the non-host plant *Nicotiana benthamiana.* Ecp20 (A) and Ecp32 (C) family members from *D. septosporum* and *F. fulva* were expressed in *N. benthamiana* using an *Agrobacterium tumefaciens-mediated* transient expression assay (ATTA) to assess their ability to elicit cell death. Representative images are shown (n = 12–24 infiltration zones), from at least three independent experiments. INF1, *Phytophthora infestans* elicitin positive cell death control; EV, empty vector negative no cell death control; Ds: *D. septosporum*; Ff: *F. fulva*. Graphs display the percentages of infiltration zones that showed cell death in response to Ecp20 (B) and Ecp32 (D) family members from *F. fulva* and *D. septosporum,* divided into three categories: strong cell death, weak cell death and no cell death. Photos were taken 7 days after infiltration.

As expected, the positive control INF1 triggered strong cell death responses and the negative control empty vector did not elicit a response. Among Ecp20 family members, FfEcp20-2 triggered strong or weak cell death responses in *N. benthamiana* in 91.7% of infiltration zones, whilst its ortholog DsEcp20-2 did not elicit any cell death responses. Both FfEcp20-3 and DsEcp20-3 caused consistent plant cell death in *N. benthamiana,* with DsEcp20-3 having 58% of infiltration zones showing strong cell death across the entire infiltration zone (Figure 5A, 5B). In *N. tabacum,* DsEcp20-3 was the only family member that triggered a cell death response (Supplementary Figure S6). Interestingly, DsEcp20-4, one of the most highly expressed secreted proteins during infection of pine by *D. septosporum* (Supplementary Table S4; Bradshaw et al. (2016)), and its ortholog from *F. fulva,* FfEcp20-4, did not induce cell death responses in either *N. benthamiana* or *N. tabacum* (Figure 5; Supplementary Figure S6).

The ability of Ecp32 family members to trigger cell death was also assessed in both *Nicotiana* species. In this case, all proteins from the Ecp32 family, apart from the Ds/FfEcp32-4 protein pair, induced some degree of cell death in both plant species. Interestingly, DsEcp32-1 consistently triggered cell death in *N. benthamiana,* while its ortholog from *F. fulva,* FfEcp32-1, only caused a strong cell death response in 3.5% of infiltration zones across all replicates (Figure 5C, 5D). In contrast, in *N. benthamiana,* FfEcp32-3 was capable of triggering consistent strong cell death responses, while DsEcp32-3 only trigged a strong response in approximately 20% of infiltration zones. The Ds/FfEcp32-5 pair triggered similar degrees of cell death responses in *N. benthamiana* (Figure 5D).

To determine whether apoplastic localization is required for members of the Ecp20 and Ecp32 families to trigger cell death, proteins that previously elicited a plant response, but lacked the PR1α signal peptide for secretion to the apoplast, were transiently expressed in *N. benthamiana.* Without the signal peptide, none of proteins tested were able to cause cell death (Supplementary Figure S7 and Supplementary Figure S8).

In cases where a cell death response was not observed at all, or where only weak cell death was observed, a Western blot analysis was performed to confirm the presence of the proteins in the tissue of the infiltrated plants (Supplementary Figure S8). All proteins that did not trigger cell death or only triggered a weak cell death response were detected via Western blot, indicating that the lack of cell death response was not due to absence of those proteins.

Taken together, these results indicate that Ecp20 and Ecp32 family members from two closely related fungal pathogens, *D. septosporum* and *F. fulva,* differ in their ability to induce a cell death response in non-host species. Furthermore, the finding that proteins without a signal peptide failed to trigger a plant cell death response, suggests that secretion into the apoplastic environment of *N. benthamiana* is essential for their cell death-eliciting activity.

### 3.5 Proteins from other *D. septosporum* Isolates and *D. pini* also Trigger Cell Death in Non-host Plants

Alleles of some *Ecp20* and *Ecp32* family genes identified from *D. pini*, and from isolates of *D. septosporum* other than NZE10, were tested for the ability of their encoded proteins to trigger cell death in *N. benthamiana*. For the proteins from *D. septosporum* NZE10 that consistently triggered the highest percentages of cell death (DsEcp20-3, DsEcp32-1 and DsEcp32-3), we tested orthologs of other *D. septosporum* alleles and found they triggered the same responses as those of *D. septosporum* NZE10 (Supplementary Figure S9). The same was true for the respective orthologs of DpEcp20-3, DpEcp32-1 and DpEcp32-3 from *D. pini*, which also elicited cell death responses in *N. benthamiana*. These results suggest conservation of cell death-eliciting activity between isolates of *D. septosporum* and between the two *Dothistroma* species for members of the Ecp20 and Ecp32 families.

### 3.6 Infiltration of Purified Ecp20 and Ecp32 Proteins from *D. septosporum* into *P. radiata* Triggers Cell Death

The pine infiltration method described in Hunziker et al. (2021) proved to be a suitable approach for testing the cell death-eliciting activity of candidate apoplastic effector proteins from pine pathogens in *P. radiata.* We previously showed that DsEcp32-3 triggered cell death in three *P. radiata* genotypes (S6, S11 and R4). Using the same method, we tested two additional proteins from the DsEcp20 family, DsEcp20-3 and DsEcp20-4, and repeated the infiltration of DsEcp32-3 as a positive cell death control (Figure 6, Supplementary Figure S10) (Hunziker et al., 2021).

**Figure 6.**
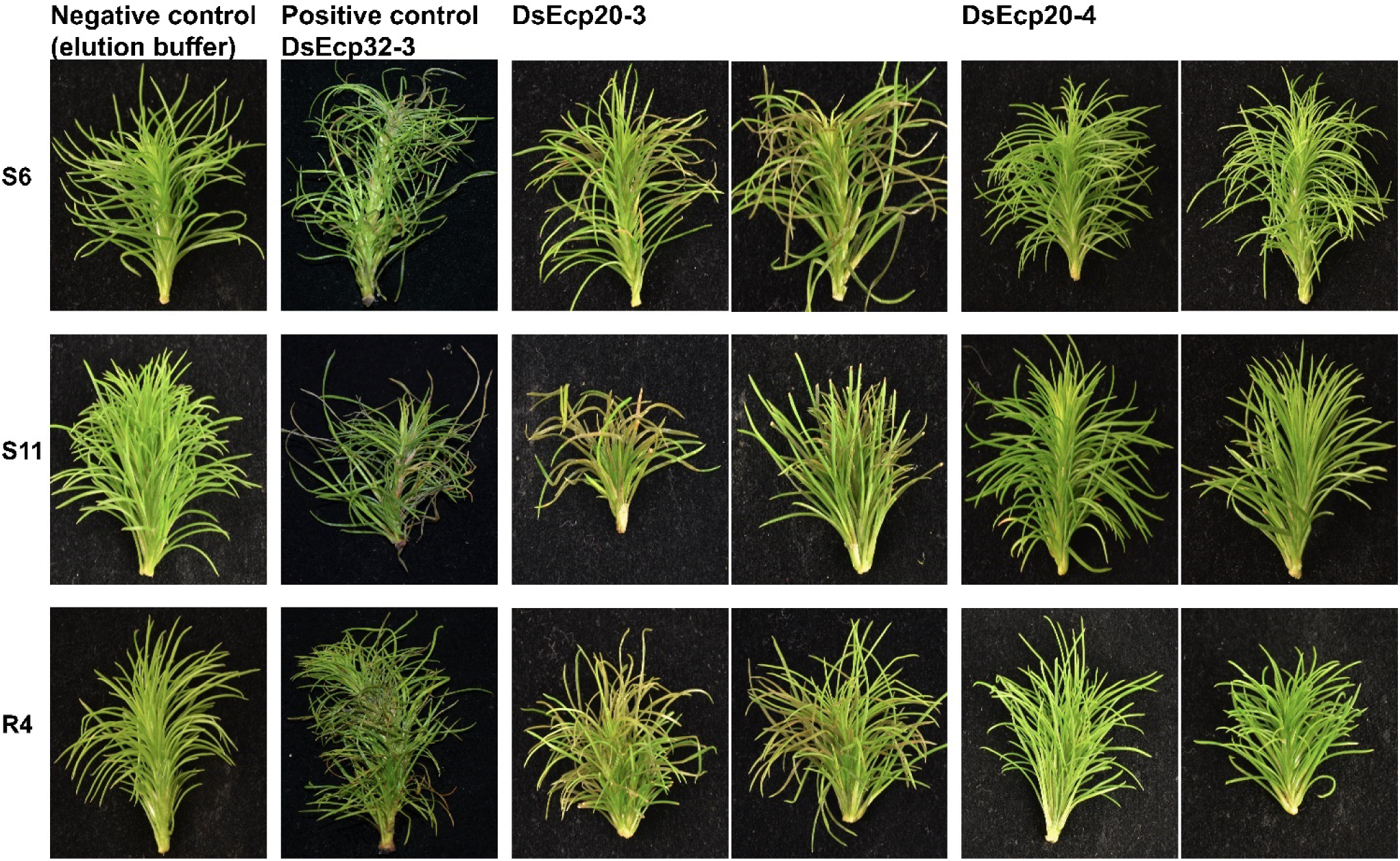
*Dothistroma septosporum* candidate effectors DsEcp20-3 and DsEcp20-4 trigger the same responses in *Pinus radiata* shoot tissues following vacuum infiltration as in non-host plants. Whole shoots of *P. radiata* derived from family seedlots that are relatively susceptible (S6 and S11) or tolerant (R4) to *D. septosporum* infection were infiltrated with purified DsEcp20-3 and DsEcp20-4 proteins produced by heterologous expression in *Pichia pastoris*. Elution buffer was used as negative control and purified DsEcp32-3 protein as a positive control. Representative photos (from 18 to 24 pine shoots for each treatment) were taken 7 days after infiltration.

The DsEcp32-3 positive control triggered cell death in all three pine genotypes, as expected (Figure 6). Infiltration of elution buffer (negative control) did not cause any damage or visible stress responses. Interestingly, both tested proteins gave the same response as they had shown in the non-host *Nicotiana* plants: DsEcp20-3 consistently elicited cell death in the same three pine genotypes used in the previous study, while DsEcp20-4 failed to elicit any response in the pine shoots.

### 3.7 DsEcp20-3-triggered Cell Death in *N. benthamiana* requires Membrane-localized Receptor-like Proteins

To determine how cell death elicitors from the Ds/FfEcp20 and 32 family were perceived by plants, we silenced genes encoding the extracellular receptor-like kinase co-receptor genes *BAK1* and *SOBIR1*, as well as leucine-rich repeat (LRR) receptor-like genes, in *N. benthamiana* (Figure 7), in conjunction with agro-infiltration, as previously described by Wang et al. (2018). As expected, agroinfiltration of positive controls NPP1, INF1 and XEG triggered consistent strong cell death responses in *N. benthamiana* leaves treated with TRV:*GFP*, while INF1- and XEG1-triggered cell death was compromised in leaves treated with *TRV:BAK1* or TRV:*SOBIR1*. We also consistently found that DsEcp20-3-triggered cell death was compromised in *N. benthamiana* leaves in which the receptorlike kinase genes *BAK1* or *SOBIR1* were silenced (Figure 7B). In addition, we found that DsEcp20-3-triggered cell death was compromised in *N. benthamiana* leaves treated with TRV VIGS constructs TRV:*T26* and TRV:*T26-2* (Wang et al., 2018) (Figure 7D). TRV:*T26* and TRV:*T26-2* both target the same two *N. benthamiana* genes but in different regions of those genes. One of these target genes encodes an LRR-type extracellular receptor-like protein (Sol Genomics Network accession number – Niben101Scf03138g01049.1) and one an extracellular LRR (Niben101Scf01834g02009.1). We further screened other members of the Ecp20 and Ecp32 families (FfEcp20-3, DsEcp32-3 and FfEcp32-3), and found that cell death triggered by FfEcp20-3 and FfEcp32-3 was also significantly compromised in *N. benthamiana* leaves treated with TRV:*BAK1*, TRV:*SOBIR1* or either of the T26 constructs (Figure 7B and D). For DsEcp32-3-triggered cell death, we did not detect any reduction on TRV:*BAK1*- or *TRV:SOBIR1-silenced* plants (data not shown). Taken together, these results suggest that the cell death responses triggered by members of the Ecp20 and Ecp32 families from *D. septosporum* and *F. fulva* in *N. benthamiana* are the result of recognition by extracellular immune receptors, and that the transduction of defense response signals following this recognition requires the extracellular co-receptors BAK1 and SOBIR1.

**Figure 7.**
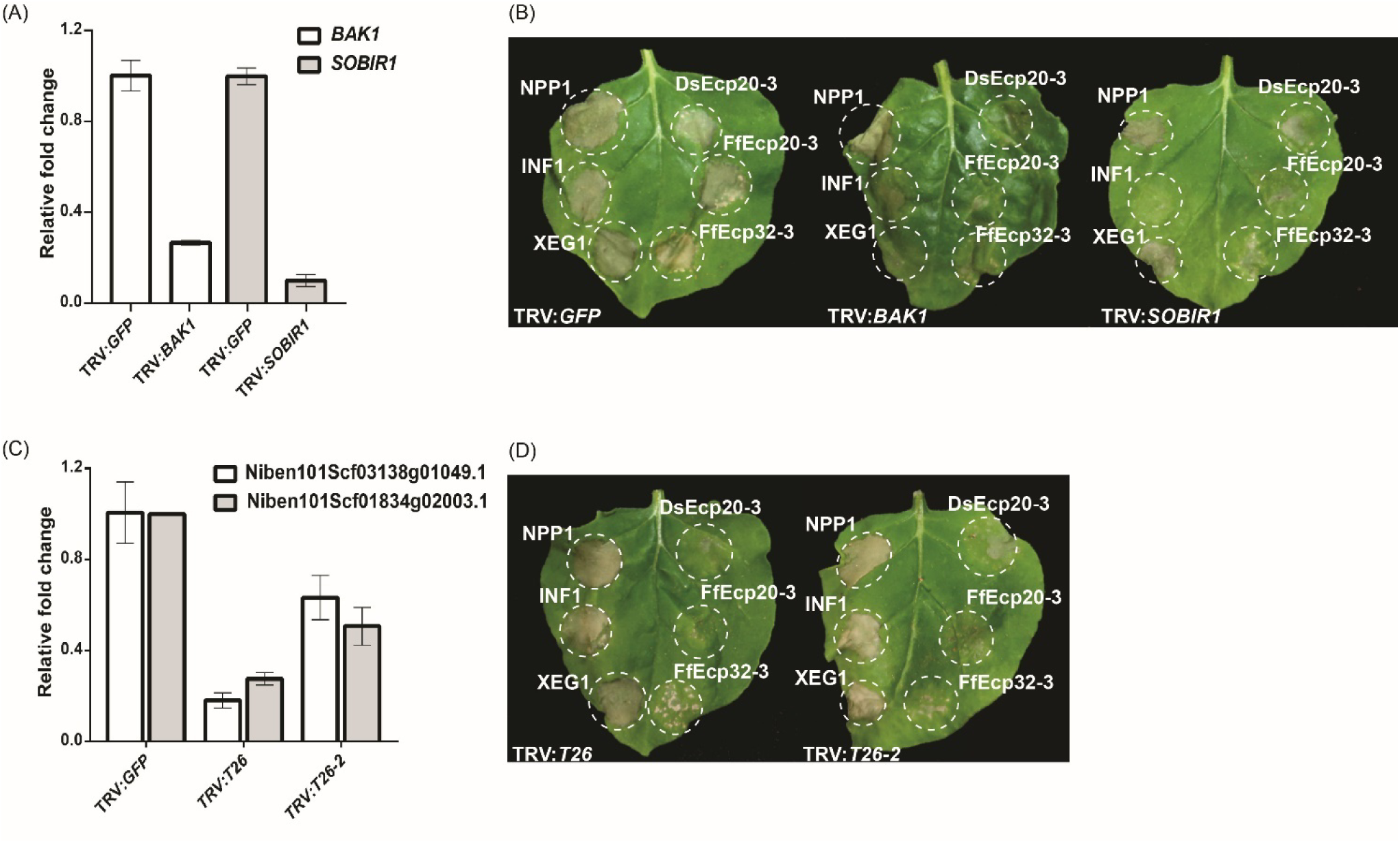
Cell death triggered by DsEcp20-3 in *Nicotiana benthamiana* is compromised in leaves silenced for different receptor-like genes. Expression levels of (A) *BAK1* and *SOBIR1* and (C) *T26* constructs in plants silenced for these genes, relative to their expression in plants silenced for the Green Fluorescent Protein (*GFP*) gene (absent in *N. benthamiana*; normalized to 1). Genes were silenced using a Virus-Induced Gene Silencing (VIGS) method based on Tobacco Rattle Virus (TRV). (C, D) Representative leaves showing cell death induced by expression of XEG1, INF1, NPP1, DsEcp20-3, FfEcp20-3 and FfEcp32-3 in *N. benthamiana* treated with specific TRV-mediated VIGS constructs as shown. Leaves (n = 8) were photographed three days after agro-infiltration (dai). Experiments were repeated twice with similar results.

## 4 Discussion

Plant-pathogenic fungi secrete a collection of virulence factors, termed effector proteins, into their hosts to promote the infection process (Lo Presti et al., 2015; Rocafort et al., 2020). While many of these effectors appear to be singletons with no obvious sequence similarity to proteins present in the effector repertoires to which they belong, others form part of families. Indeed, in *Venturia inaequalis*, the biotrophic Dothideomycete pathogen responsible for scab disease of apple, more than half of the predicted effector repertoire is made up of protein families based on sequence similarity alone (Deng et al., 2017; Rocafort et al., 2022). Likewise, in *Ciborinia camelliae,* the necrotrophic Leotiomycete pathogen responsible for flower blight disease of camellias, approximately one tenth of the predicted secretome belongs to a single massively expanded candidate effector family (Denton-Giles et al., 2020). Notably, effector families can also be associated with avirulence. For example, in the hemibiotrophic Sordariomycete pathogen *Magnaporthe oryzae* (rice blast disease), the biotrophic Leotiomycete pathogen *Blumeria graminis* (cereal powdery mildew disease), as well as the biotrophic Pucciniomycete pathogen *Melampsora lini* (flax rust disease) and the hemibiotrophic Dothideomycete pathogen *Leptosphaeria maculans* (stem canker disease of brassica), certain avirulence effectors have been found to form part of protein families (Anderson et al., 2016; Catanzariti et al., 2006; Dodds et al., 2004; Kang et al., 1995; Khang et al., 2008; Praz et al., 2017; Spanu, 2017; Talbi et al., 2022).

The emergence of sequence-related effector families in fungi is often the result of duplication events from an ancestral gene or from other family members and is typically followed by sequence diversification through an accumulation of indels and synonymous/non-synonymous substitutions and/or recombination events between different family members (Denton-Giles et al., 2020; Franceschetti et al., 2017; Pendleton et al., 2014). Such sequence diversification is anticipated to have roles in, for example, the diversification of effector functions, as well as the evasion of host immunity. In some cases, this diversification can result in an almost complete loss of sequence similarity between family members, with similarity only retained at the protein tertiary structure level, as has recently been shown for several effector families of plant-pathogenic fungi using the groundbreaking *ab initio* protein structure prediction tool, AlphaFold2 (Rocafort et al., 2022; Seong & Krasileva, 2022). However, in many cases, sequence identity between family members is maintained, even across different fungal species. These cross-species effector families are of high interest, as their sequence conservation may indicate an important and/or conserved (i.e., core) role in fungal virulence (Stergiopoulos et al., 2010).

In this study, we identified two candidate effector protein families, Ecp20 and Ecp32, which are highly conserved at the primary sequence and tertiary structure levels between the two closely related Dothideomycete fungal pathogens, *D. septosporum* and *F. fulva.* Despite their close phylogenetic relationship, these two pathogens have very different hosts, with *D. septosporum* causing disease on pine (a gymnosperm), and *F. fulva* causing disease on tomato (an angiosperm) (Drenkhan et al., 2016; Thomma et al., 2005). An investigation of pre-existing gene expression data revealed that both protein families have members encoded by genes that are highly expressed *in planta,* while others are only moderately or lowly expressed. Such a finding is consistent with what was recently observed in *V. inaequalis,* where gene family members are expressed at different levels during apple infection (Rocafort et al., 2022).

Both the Ecp20 and Ecp32 families were found to have members that differed in their ability to trigger cell death in the non-host species *N. benthamiana* and *N. tabacum.* The differential ability of these family members to trigger a cell death response does not appear to be restricted to proteins from *D. septosporum* and *F. fulva.* Indeed, it was recently determined that an ortholog of Ds/FfEcp20-4 from the hemibiotrophic Dothideomycete pathogen *Zymoseptoria tritici* (septoria leaf blotch disease of wheat), named Zt4 (JGI protein ID 104697), triggers cell death in *N. benthamiana* (Kettles et al., 2017). This is in contrast to Ds/FfEcp20-4, where, in our study, no cell death response was observed. Similarly, a homolog of Ecp32-1 from *Z. tritici* (JGI protein ID 105896; no reciprocal ortholog in *D. septosporum* or *F. fulva)* was not able to induce a cell death response in *N. benthamiana* (Kettles et al., 2017) whereas in our study, DsEcp32-1 and FfEcp32-1 consistently and infrequently triggered cell death, respectively. Of course, it remains possible that some of the family members triggered other plant responses, such as changes in host gene expression or the accumulation of reactive oxygen species (ROS), which were not measured in our assays. In any case, variation in the ability of family members to elicit plant cell death is not restricted to the Ecp20 and Ecp32 families, with variation also observed in, for example, the cell death-inducing capacity of CcSSP effector candidate family members from *C. camelliae* in camellia petals (Denton-Giles et al., 2020).

BLASTp searches demonstrated that, like those shown above for *Z. tritici,* homologs of the Ecp20 and Ecp32 family members are present in other fungal species, with many of these species being plant pathogens. Interestingly, as observed in *D. septosporum* and *F. fulva,* these homologs were also frequently found to exist as protein families. Taken together, it is therefore likely that the Ecp20 and Ecp32 proteins are core effectors, possibly contributing to conserved virulence functions in their respective plant hosts. In addition to searching for homologs in other plant pathogens, we also screened the genomes of other *D. septosporum* isolates and of *D. pini,* the other causal species of DNB, and found that whilst Ecp20 and Ecp32 family members are polymorphic across other *D. septosporum* isolates, all family members are present in these isolates and also in *D. pini,* with the exception of Ecp32-5. There was also conservation of cell death-eliciting function between isolates of *D. septosporum* and between the two *Dothistroma* species, since the respective orthologs showed the same responses as those from the NZE10 strain, also triggering cell death in *N. benthamiana.* Whether the identified sequence variation contributes to virulence function of these proteins outside of cell death elicitation remains to be determined.

As the primary amino acid sequence of the Ecp20 and Ecp32 family members provided no insights into their potential virulence function, we predicted the tertiary structures of these proteins using the AlphaFold2 tool and compared them to proteins of characterized structure and/or function. Based on this analysis, all eight *D. septosporum* and *F. fulva* Ecp20 family members were predicted to possess structural similarity to PevD1, an effector protein with an Alt a 1 (β-barrel) fold from the hemibiotrophic Sordariomycete pathogen *Verticillium dahliae* (Chruszcz et al., 2012; Han et al., 2012). Interestingly, the Alt a 1 fold has also been found in MoHrip1, a plasma membrane-binding virulence factor from *M. oryzae* that is highly expressed during rice infection (Zhang et al., 2017). Just like FfEcp20-2, FfEcp20-3 and DsEcp20-3, the PevD1 and MoHrip1 proteins induce cell death responses in *Nicotiana* species (Han et al., 2012; Zhang et al., 2017), suggesting that this cell death-eliciting ability could be a conserved function of these proteins. It should be noted, however, that proteins with an Alt a 1 fold, including PevD1, have also been shown to interact with and inhibit the enzymatic activity of pathogenesis-related 5 (PR5) proteins from different host plants (Gómez-Casado et al., 2014; Kumar & Mukherjee, 2020; Zhang et al., 2019). Whether the Ecp20 proteins from *D. septosporum* and *F. fulva* perform a similar role in pathogen virulence remains to be explored.

In contrast to members of the Ecp20 family, the Ecp32 proteins were predicted to be structurally similar to proteins from diverse organisms with a β-trefoil fold. Proteins with a β-trefoil fold have the ability to recognize different ligands, such as DNA, proteins and carbohydrates, and so can be involved in different processes and interactions between organisms (Žurga et al., 2015). Among the β-trefoil fold proteins, the most abundant members are lectins, carbohydrate-binding proteins and hydrolase inhibitors, such as protease inhibitors (Renko et al., 2012). Lectins are proteins that specifically bind carbohydrates and, in fungi, are associated with plant-pathogen interactions, fungal growth, fruiting body development and, due to their toxicity, defense against parasitic or predatory organisms (Sharon, 2007; Varrot et al., 2013). Lectins from different mushroom species, for example, are known to be toxic to organisms such as *Caenorhabditis elegans* and *Drosophila melanogaster* (Juillot et al., 2016; Schubert et al., 2012; Wohlschlager et al., 2011). These proteins bind to the plasma membrane and, in doing so, induce cell death. Cyt toxins from *Bacillus thuringiensis,* for instance, are insecticidal proteins with a β-trefoil fold that are known to interact with host membrane lipids (Soberón et al., 2013). Even though the mechanism of lectin toxicity is not known, it is hypothesized that this toxicity may be the direct result of host membrane damage. Taken together, this could indicate a possible role Ecp32 proteins in triggering plant cell death, at least in part, by interacting with and disrupting the host membrane.

As mentioned above, protease inhibitors can also adopt a β-trefoil fold, and in our study, it was found that most Ds/FfEcp32 proteins had a predicted tertiary structure similar to that of a plant trypsin inhibitor. Of interest, fungal pathogens frequently target host proteases involved in plant defense by secreting effectors that serve as protease inhibitors, with well characterized examples including Pit2 from the biotrophic Ustilaginomycete *Ustilago maydis* (corn smut disease) and Avr2 from *F. fulva* (Mueller et al., 2013; Tian et al., 2007; van Esse et al., 2008). While it is not yet known whether Pit2 or Avr2 adopt a β-trefoil fold, another possible function for the Ecp32 proteins could therefore involve the inhibition of plant proteases during host colonization.

The fact that Ecp20 and Ecp32 family members of *D. septosporum, F. fulva* and *Z. tritici* are likely apoplastic (Hunziker et al., 2021; Kettles et al., 2017; Mesarich et al., 2018), and that members of both families across multiple fungal species trigger cell death in *Nicotiana* species (with this activity dependent on their secretion to the apoplast), may suggest that each family presents a conserved epitope that is recognized by an extracellular immune receptor in these non-host plants. In support of recognition by an extracellular immune receptor, silencing of the extracellular co-receptors *NbBAK1* or *NbSOBIR1* in *N. benthamiana,* which are required to transduce defense response signals following apoplastic effector recognition (Liebrand et al., 2013; Liebrand et al., 2014), compromised Ds/FfEcp20-3-dependent cell death. Notably, a compromised cell death response was also observed for FfEcp32-3, but not DsEcp32-3. One possibility here is that the expression of *NbBAK1* and/or *NbSOBIR1* was not reduced to a level sufficient for the cell death response triggered by DsEcp32-3 to be compromised. Alternatively, given its predicted structural similarity to lectins, other processes outside of immune receptor recognition, such as cell membrane perturbation, could be involved in cell death elicitation by DsEcp32-3.

In line with the cell death response observed in the non-host species, we previously determined that DsEcp32-3 can trigger cell death in pine and suggested that this protein might be a virulence factor in the necrotrophic stage of DNB that helps to destroy needle tissue (Hunziker et al., 2021). With this in mind, we decided to investigate whether two (purified) *D. septosporum* proteins from the Ecp20 family could also induce a cell death response in pine. Both proteins, DsEcp20-3 and DsEcp20-4, were of interest due to their high expression in the Mid and Late (necrotrophic) stages of *D. septosporum* infection in *P. radiata* (Bradshaw et al., 2016). The finding that DsEcp20-3 and DsEcp20-4 triggered different cell death responses in non-host plants, *N. benthamiana* and *N. tabacum,* also made them good candidates for this assay. Interestingly, DsEcp20-3, like DsEcp32-3, was able to trigger a cell death response in all pine genotypes tested. Taken together, these results suggest that the DsEcp20-3 and DsEcp32-3 proteins could also by recognized by an immune receptor (possibly conserved with *N. benthamiana)* or, as described above, participate in membrane perturbation in *P. radiata.* Consistent with what was observed in the non-host plants, DsEcp20-4 was unable to elicit a cell death response in the pine shoots. This may suggest that it does not have the conserved epitope required to elicit cell death following recognition by an immune receptor. To identify the regions or amino acid residues involved in cell death induction, and thus potential epitopes for recognition by cognate immune receptors, region swaps between cell death-eliciting and non-eliciting members of the Ecp20 and Ecp32 families, based on information gained from the prediction of surface-exposed residues by AlphaFold2, can now be performed in *Nicotiana* species or pine, similar to that described for the Avr4 effector protein of *D. septosporum* and *F. fulva* in *N. benthamiana* (Mesarich et al., 2016). This, in turn, might help to explain the differences in cell death-inducing capacity between, for example, CfEcp32-1 and DsEcp32-1, as well as DsEcp20-3 and DsEcp20-4.

As it is well known that plant cell death is a strategy exploited by necrotrophic and hemibiotrophic pathogens to acquire nutrients from their hosts during their necrotrophic stage or to facilitate a biotrophic-necrotrophic switch (Shao et al., 2021), one possibility is that the DsEcp20-3 and DsEcp32-3 proteins from *D. septosporum* are involved in these processes. In addition to their cell death-eliciting activities, support for this hypothesis is provided by the fact that the expression of both *DsEcp20-3* and *DsEcp32-3* is highly induced during the Mid and Late stages of pine infection by *D. septosporum*. Another possibility is that DsEcp20-3 may be acting as an Avr factor of *D. septosporum* recognized by the host, similar to that already proposed for the core effector DsEcp2-1 (Guo et al., 2020), while other family members, such as DsEcp20-4, which is also highly expressed during pine infection, might be virulence factors that interact with pine PR5 proteins to suppress host defenses. One way to approach these research questions is to generate mutants of *D. septosporum* deleted for a *DsEcp20* or *DsEcp32* gene (e.g., using CRISPR/Cas9 technology) to determine what impact their deletion or disruption has on necrotrophic infection by this fungus. The problem with this method is that because these genes form part of multigene families, functional redundancy may mask the role of the individual effectors (i.e. the deletion mutants may not show a reduction in virulence). However, it may be possible to create strains in which genes corresponding to several or all family members are deleted to assess whether fungal virulence, particularly in the necrotrophic infection stage, is compromised. Recently, we reported the first successful application of CRISPR/Cas9 genome editing to *D. septosporum* by disrupting a gene encoding a secreted protein that elicited cell death in *N. benthamiana* and pine (McCarthy et al., 2022), which will ultimately help with creating such strains.

In conclusion, our study describes two new candidate effector families with cell death-eliciting activity from *D. septosporum* and *F. fulva* and provides evidence to suggest that members of these families are recognized as invasion patterns by immune receptors of phylogenetically distinct plant species (angiosperms versus a gymnosperm). By characterizing the virulence (and potential avirulence) functions of such ‘core’ effectors, the knowledge gained could ultimately be used in durable disease resistance programs active against these two, and other, important plant pathogens.

## 6 Author Contributions

R.B., R.M., Y.W. and C.M. conceived and guided the study. M.T. and Z.C. designed and performed experiments and analyzed data. M.T., C.M. and R.B. led manuscript writing. All authors contributed to the manuscript.

## 8 Acknowledgments

We thank Melissa Guo (Massey University, Palmerston North, New Zealand) and Lukas Hunziker (Curtin University, Perth, Australia) for input on experiments. Trevor S. Loo (Massey University, Palmerston North, New Zealand) is acknowledged for his advice on performing the IMAC, and Keiko Gough (Scion, Rotorua, New Zealand) for preparing the pine material.

