## Supplementary Figures for "Characterization of two conserved cell death elicitor families from the Dothideomycete fungal pathogens *Dothistroma septosporum* and *Fulvia fulva* (syn. *Cladosporium fulvum*)"

(A)

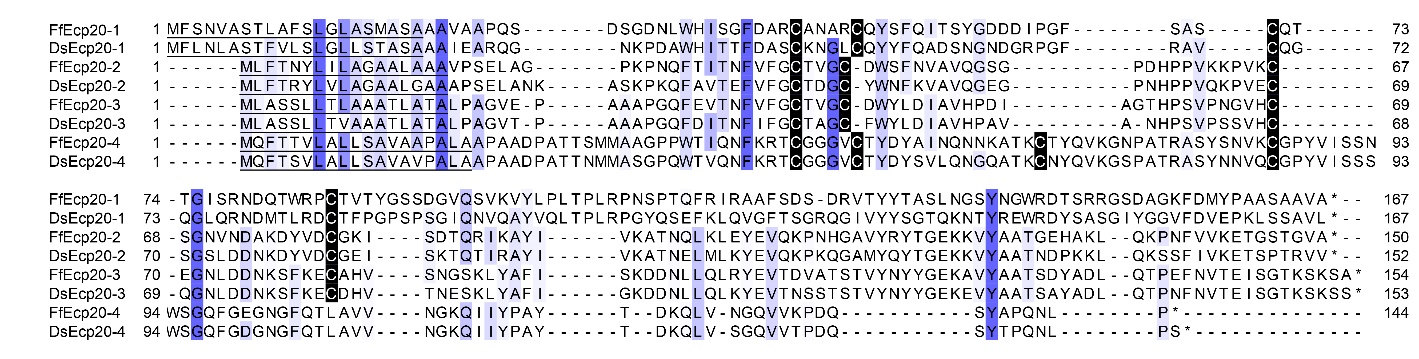

(B)

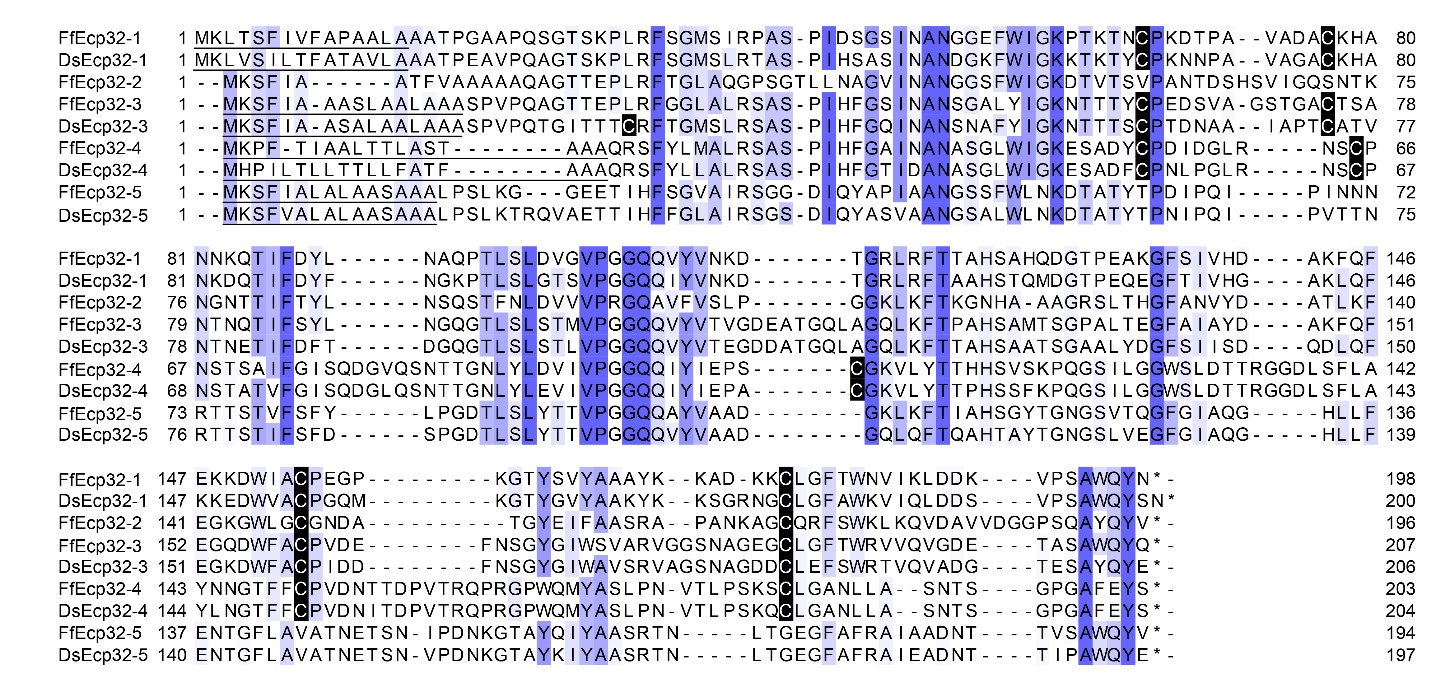

**Supplementary Figure S1.** Protein alignment of *Dothistroma septosporum* and *Fulvia fulva* Ecp20 (A) and Ecp32 (B) family members**. The alignment was generated using Clustal** Ω (Sievers et al., 2011)**, with blue** shading indicating the degree of amino acid sequence conservation. Conserved cysteine residues are highlighted in black and predicted signal peptides are underlined.

(A)

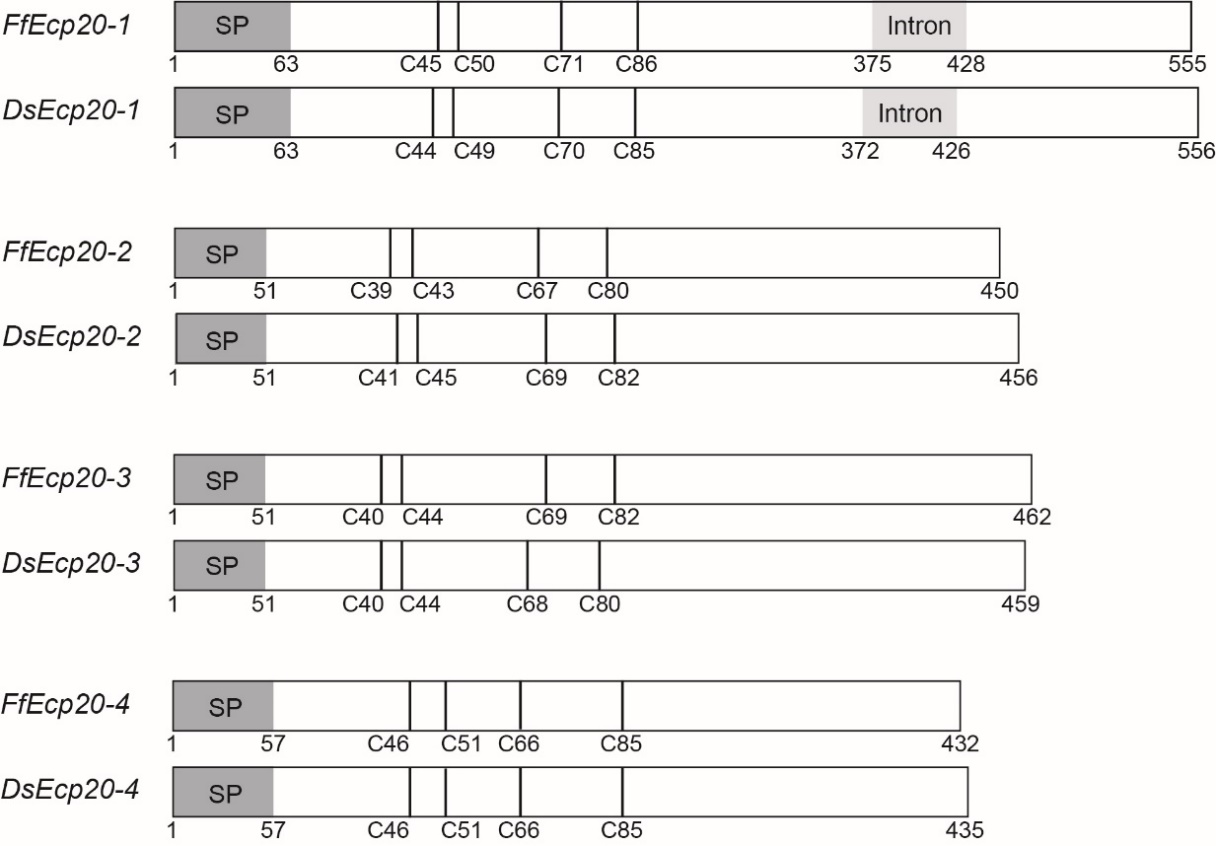

(B)

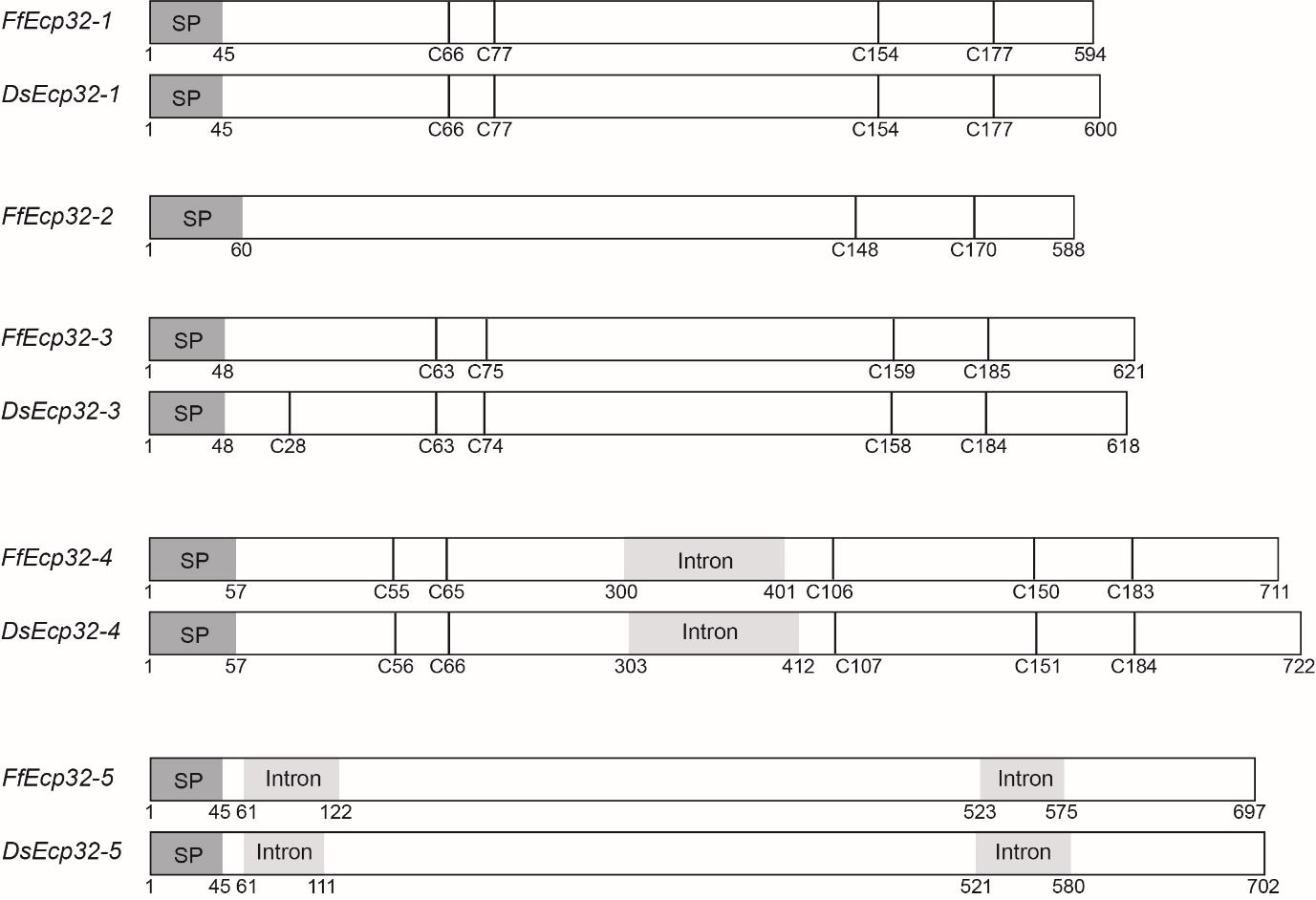

**Supplementary Figure S2.** Schematic representation of *Ecp20* (A) and *Ecp32* (B) gene family members. Dark grey boxes indicate the signal peptide (SP) position and light grey boxes indicate introns. Vertical black lines indicate cysteine residues, with the respective amino acid position shown under each gene.

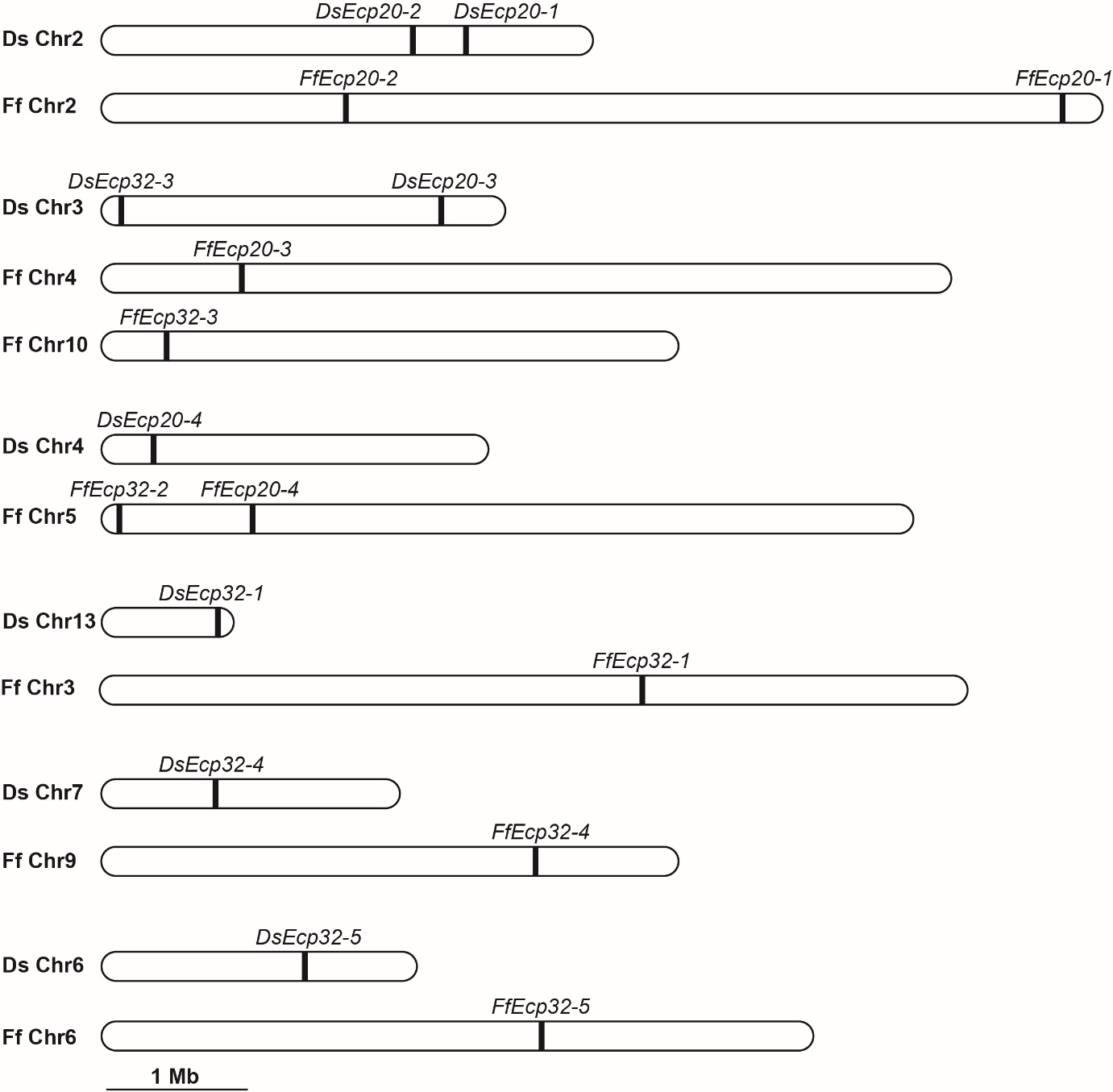

**Supplementary Figure S3.** Chromosome locations of *Dothistroma septosporum* NZE10 (Ds) and *Fulvia fulva* Race 5 (Ff) *Ecp20* and *Ecp32* genes. Matching chromosomes are grouped together, with Ff Chr4 and Ff Chr10 both matching with Ds Chr3 from *D. septosporum* (Zaccaron et al., 2022).

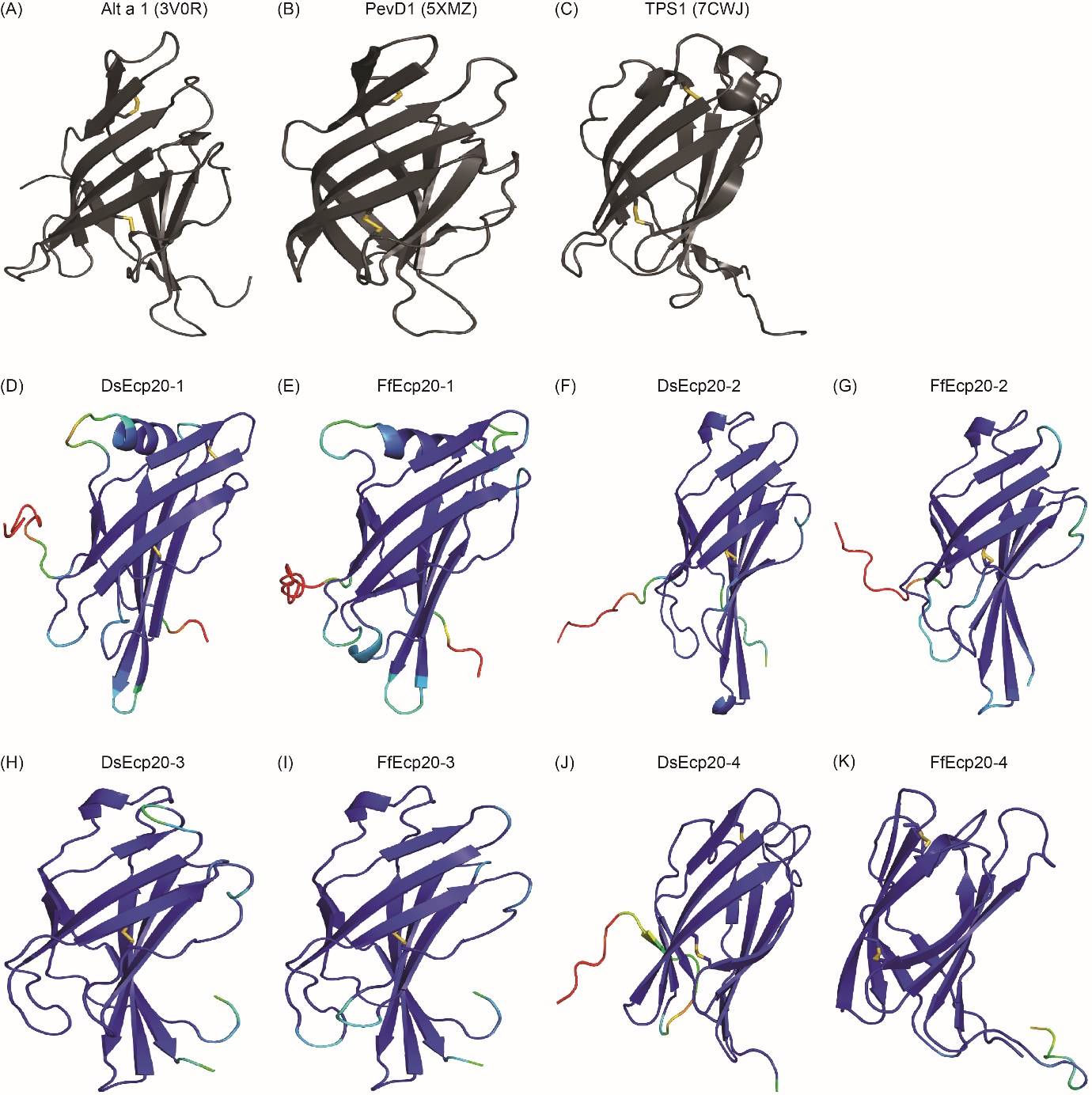

**Supplementary Figure S4. Predicted protein tertiary structures of Ecp20 family members** **from *Dothistroma septosporum* and *Fulvia fulva*. (A) Solved crystal structure of Alt a 1 from *Alternaria alternata* (RCSB protein data bank [PBD] ID: 3V0R) (Chruszcz et al., 2012). (B) Solved crystal structure of PevD1 from *Verticillium dahliae* (RCSB PBD ID: 5XMZ) (Zhou et al., 2017), (C) Solved crystal structure of TSP1 from *Trichoderma virens* (RCSB PBD ID: 7CWJ) (Gupta et al., 2021). (D-K) Predicted structure of (D) DsEcp20-1. (E) FfEcp20-1. (F) DsEcp20-2. (G) FfEcp20-2. (H) DsEcp20-3. (I) FfEcp20-3. (J) DsEcp20-4. (K) FfEcp20-4. Disulphide bonds are shown as yellow sticks. Protein structures were predicted with AlphaFold2 and rendered in PyMol v2.5 (DeLano, 2002; Jumper et al., 2021; Mirdita et al., 2022). Structures D–K were colored in PyMol according to their AlphaFold2 pLDDT score:** dark blue for highly confident predicted regions, light blue and green for regions of low confidence, and red for very low confidence regions.

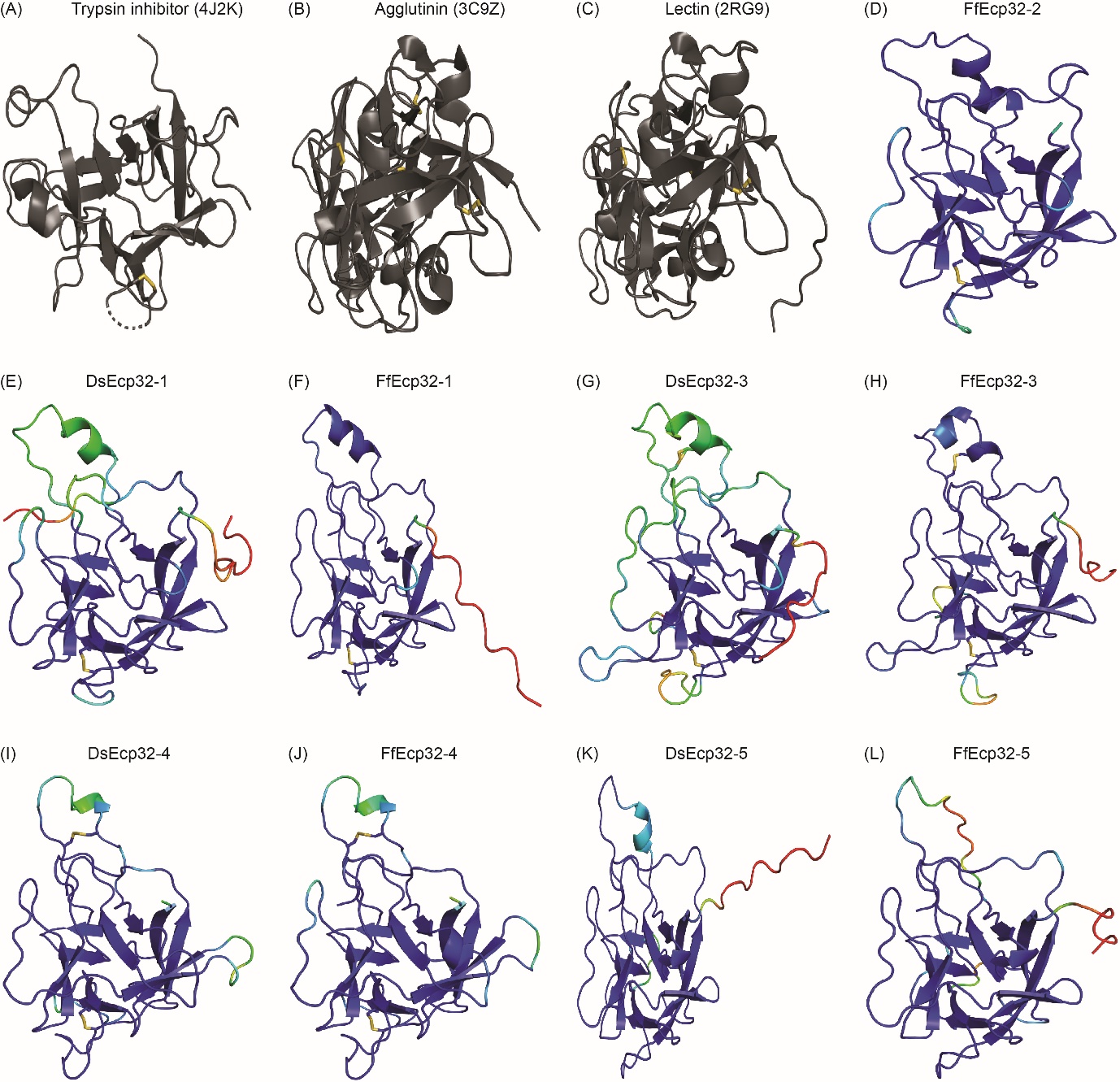

**Supplementary Figure S5. Predicted protein tertiary structures of Ecp32 family members from *Dothistroma septosporum* and *Fulvia fulva*. (A) Solved crystal structure of trypsin inhibitor from *Enterolobium contortisiliquum* (RCSB protein data bank [PBD] ID: 4J2K) (Zhou et al., 2013). (B) Solved crystal structure of agglutinin from *Sambucus nigra* (RCSB PBD ID: 3C9Z) (Maveyraud et al., 2009). (C) Solved crystal structure of lectin from *Viscum album* (RCSB PBD ID: 2RG9) (**Karpechenko et al., unpublished**). (D-L) Predicted structures of (D) FfEcp32-2. (E) DsEcp32-1. (F) FfEcp32-1. (G) DsEcp32-3. (H) FfEcp32-3.** (I) **DsEcp32-4.** (J) **FfEcp32-4. (K) DsEcp32-5. (L) FfEcp32-5. Disulphide bonds are shown as yellow sticks. Protein structures were predicted with AlphaFold2 and rendered in PyMol v2.5 (DeLano, 2002; Jumper et al., 2021; Mirdita et al., 2022). Structures D–L were colored in PyMol according to their AplhaFold2 pLDDT score:** dark blue for highly confident predicted regions, light blue and green for regions of low confidence, and red for very low confidence regions.

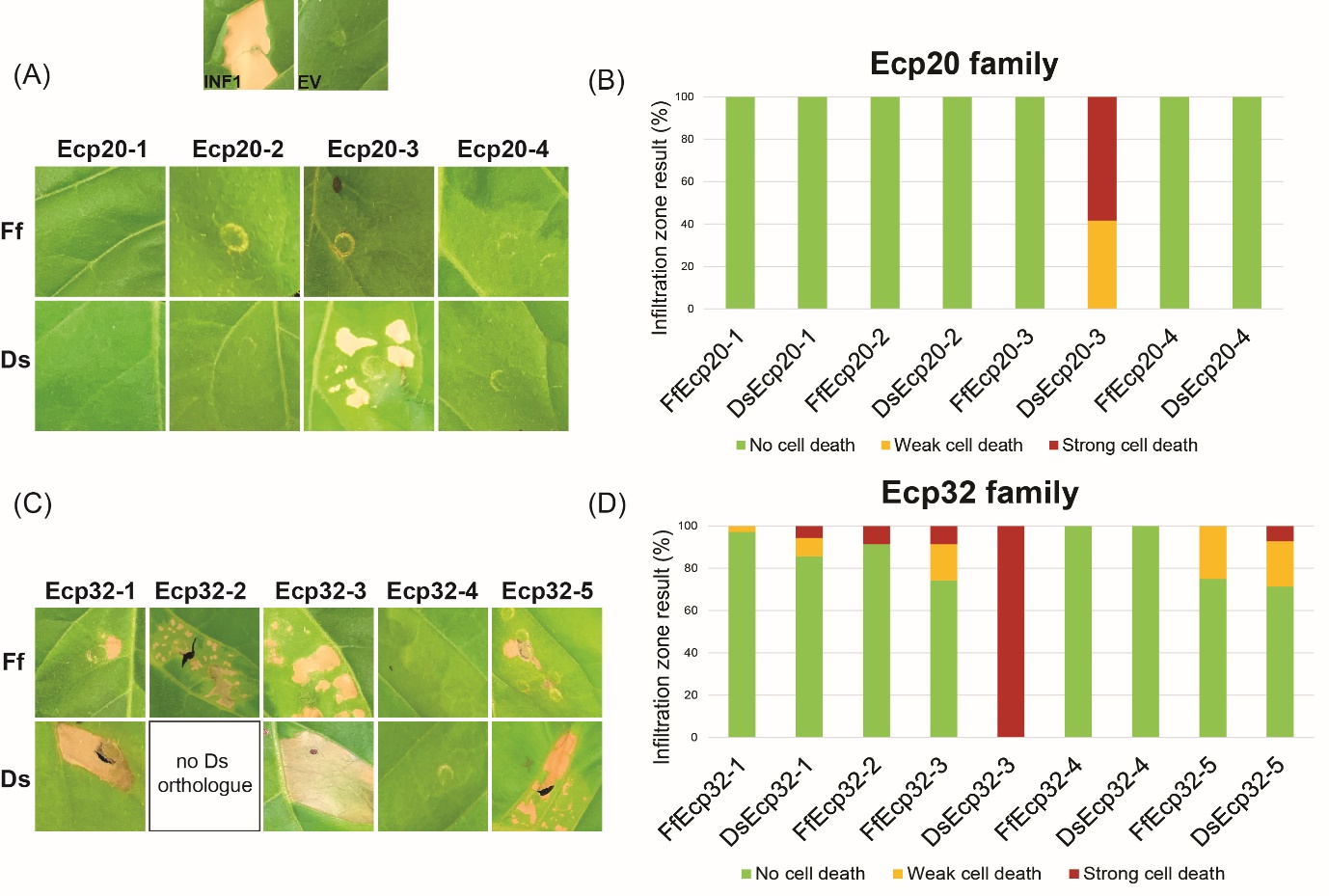

**Supplementary Figure S6.** Ecp20 and Ecp32 proteins from *Dothistroma septosporum* NZE10 and *Fulvia fulva* 0WU trigger cell death responses in the non-host plant *Nicotiana tabacum*. Ecp20 (A) and Ecp32 (C) family members from *D. septosporum* and *F. fulva* were expressed in *N. tabacum* using an *Agrobacterium tumefaciens*-mediated transient expression assay (ATTA) to assess their ability to elicit cell death. Representative images are shown (n = 12–24 inﬁltration zones), from at least three independent experiments. INF1, *Phytophthora infestans* elicitin positive cell death control; EV, empty vector negative no cell death control; Ds: *D. septosporum*; Ff: *F. fulva*. Graphs display the percentage of infiltration zones that showed cell death in response to Ecp20 (B) and Ecp32 (D) family members from *F. fulva* and *D. septosporum*, divided into three categories: strong cell death, weak cell death and no cell death. Photos were taken 7 days after infiltration.

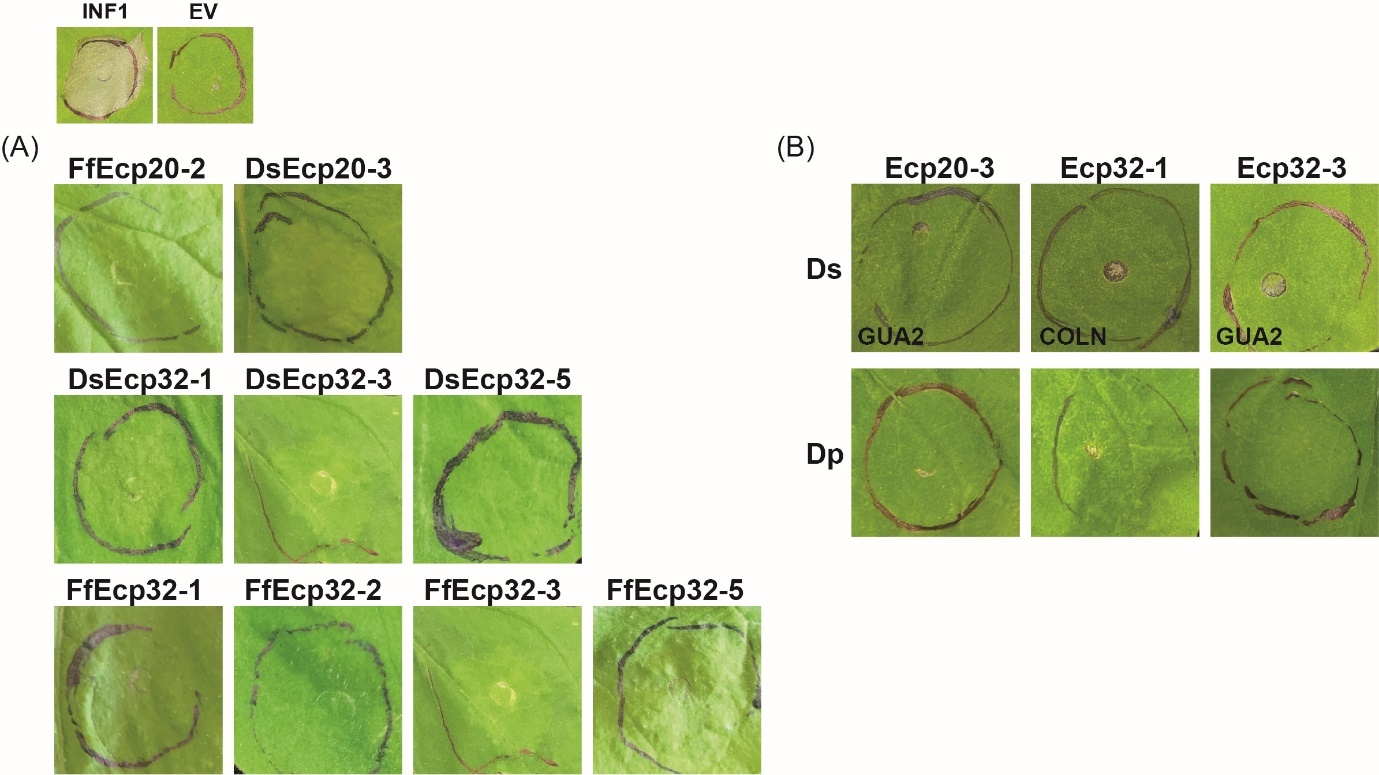

**Supplementary Figure S7.** Apoplastic localization is required for cell death elicitation by members of the Ecp20 and Ecp32 protein families from *Dothistroma septosporum* and *Fulvia fulva* in *Nicotiana benthamiana*. Ecp20 and Ecp32 proteins from *D. septosporum* NZE10 and *F. fulva* 0WU (A), and from other *D. septosporum* strains and *Dothistroma pini* (B), were expressed without a signal peptide in *N. benthamiana* using an *Agrobacterium tumefaciens*-mediated transient expression assay (ATTA) to assess their ability to elicit cell death. INF1, *Phytophthora infestans* elicitin positive cell death control; EV, empty vector negative no cell death control; Ds: *D. septosporum*; Ff: *F. fulva*. Photos were taken 7 days after infiltration.

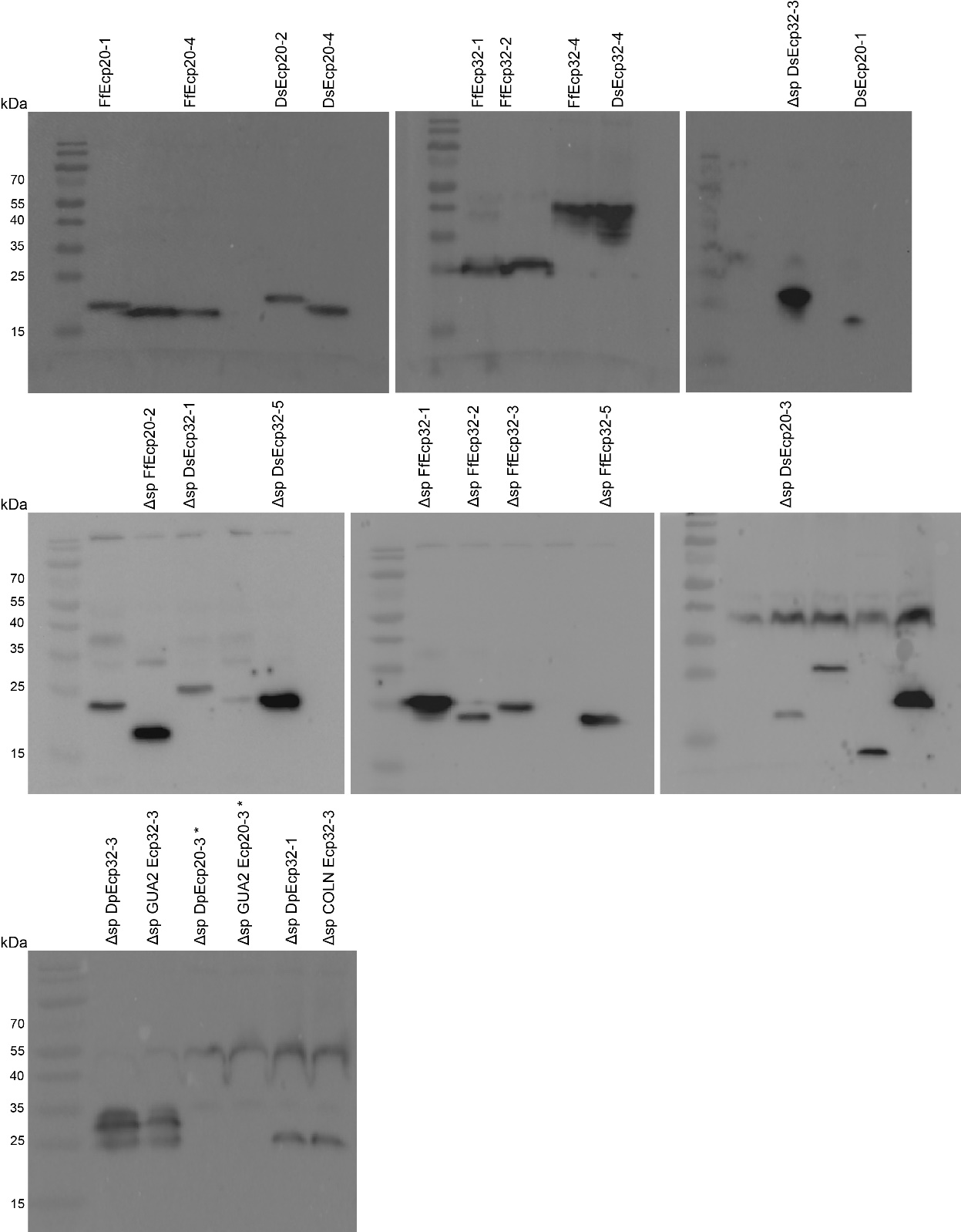

**Supplementary Figure S8.** Western blots of *Fulvia fulva*, *Dothistroma septosporum* and *Dothistroma pini* Ecp20 and Ecp32 proteins. Western blot detection of Ecp20 and Ecp32 proteins that did not trigger a cell death response or triggered only a weak cell death response when expressed in *Nicotiana benthamiana* and from Ecp20 and Ecp32 proteins that previously triggered a cell death response in *N. benthamiana*, but that now lack the PR1α signal peptide (Δsp). Expected sizes (kDa): FfEcp20-1: 17.3; FfEcp20-4: 16.2; DsEcp20-2: 17.1; DsEcp20-4: 16.2; FfEcp32-1: 22.6; FfEcp32-2: 21.6; FfEcp32-4: 22.5; DsEcp32-4: 22.6; DsEcp20-1: 19; Δsp DsEcp32-3: 22.8; Δsp FfEcp20-2: 17.2; Δsp DsEcp32-1: 22.8; Δsp DsEcp32-5: 22.2; Δsp FfEcp32-1: 22.4; Δsp FfEcp32-2: 21.4; Δsp FfEcp32-3: 22.6; Δsp FfEcp32-5: 21.7; Δsp DsEcp20-3: 17.5; Δsp DpEcp32-3: 22.8; Δsp GUA2 Ecp32-3: 22.8; Δsp DpEcp32-1: 22.8; Δsp COLN Ecp32-1: 22.6. *Δsp DpEcp20-3 and Δsp GUA2 Ecp20-3 proteins could not be detected. Anti-FLAG antibody was used for immuno-detection. Marker on the far left of membranes was the PageRuler™ Prestained Protein Ladder (ThermoScientific). Bands that are not accompanied by a label do not relate to this study.

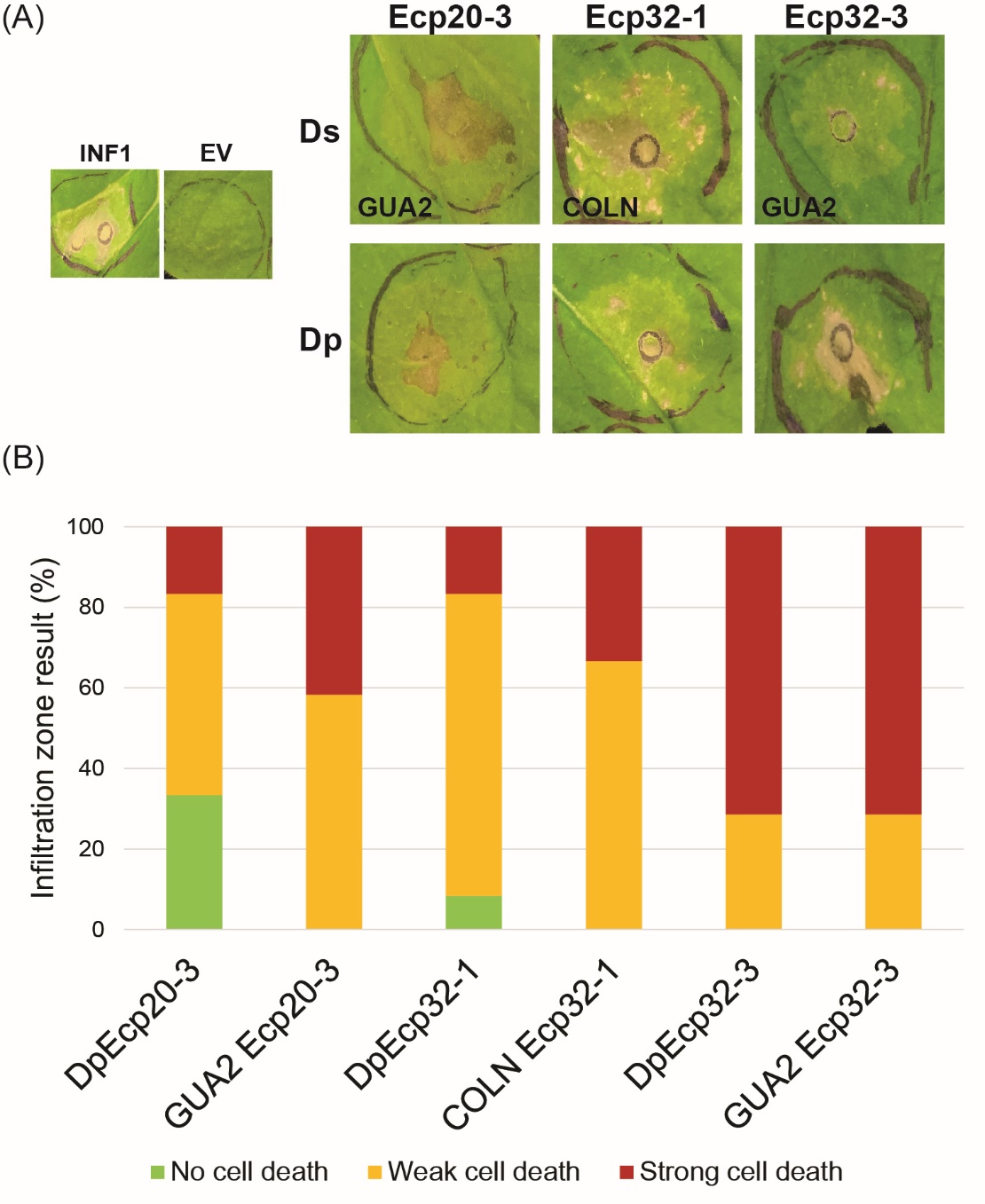

**Supplementary Figure S9.** Ecp20 and Ecp32 proteins from other *Dothistroma septosporum* isolates and *Dothistroma pini* trigger cell death in non-host plant *Nicotiana benthamiana*. (A) DsEcp20-3 and DsEcp32-3 from the Guatemalan *D. septosporum* isolate (GUA2), DsEcp32-1 from the Colombian *D. septosporum* isolate (COLN), together with their orthologues from *D. pini*, were expressed in *N. benthamiana* using an *Agrobacterium tumefaciens*-mediated transient expression assay (ATTA) to assess their ability to elicit cell death. Representative images are shown (n = 12–24 inﬁltration zones), from at least three independent experiments. INF1, *Phytophthora infestans* elicitin positive cell death control; EV, empty vector negative no-cell death control; Ds: *D. septosporum*; Dp: *D. pini*. Graphs display the percentage of infiltration zones that showed cell death in response to the respective Ecp20 and Ecp32 family members tested from *D. pini* and the most polymorphic *D. septosporum* isolates, divided into three categories: strong cell death, weak cell death and no cell death. Photos were taken 7 days after infiltration.

(A)

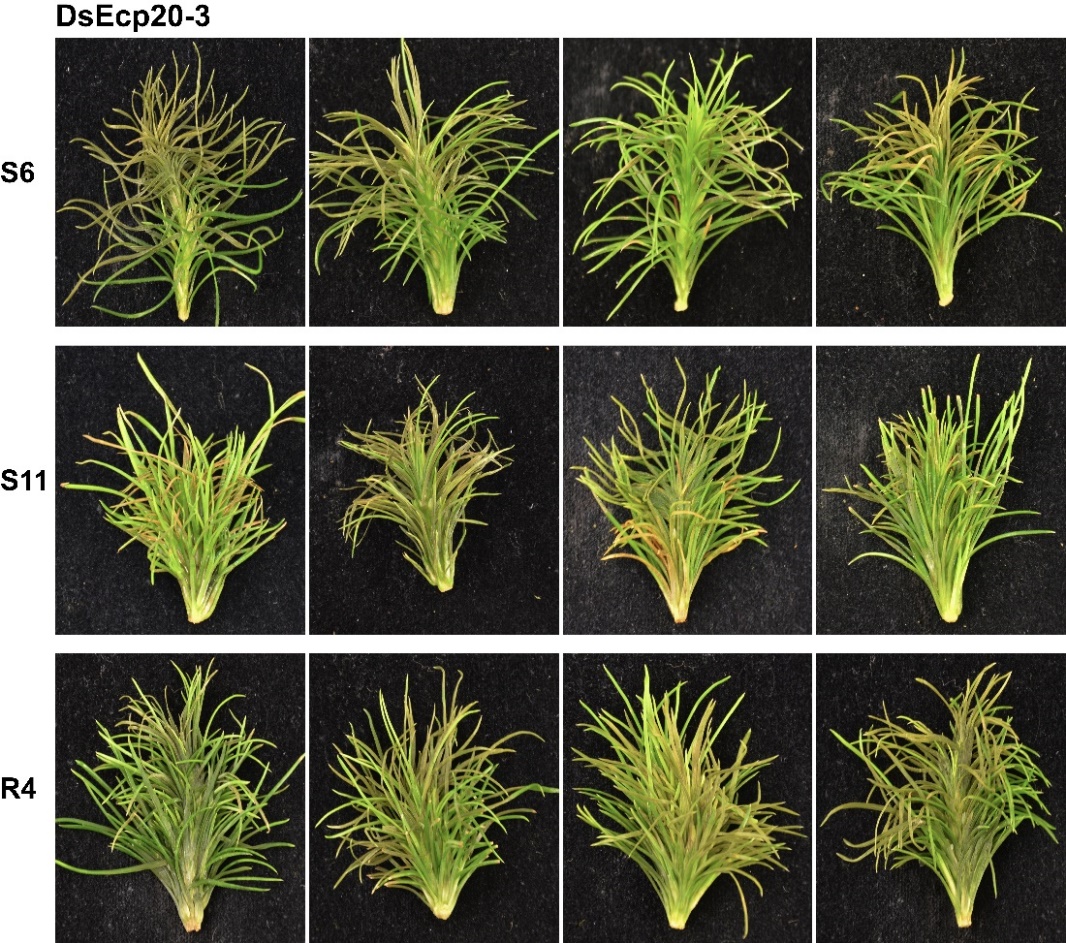

(B)

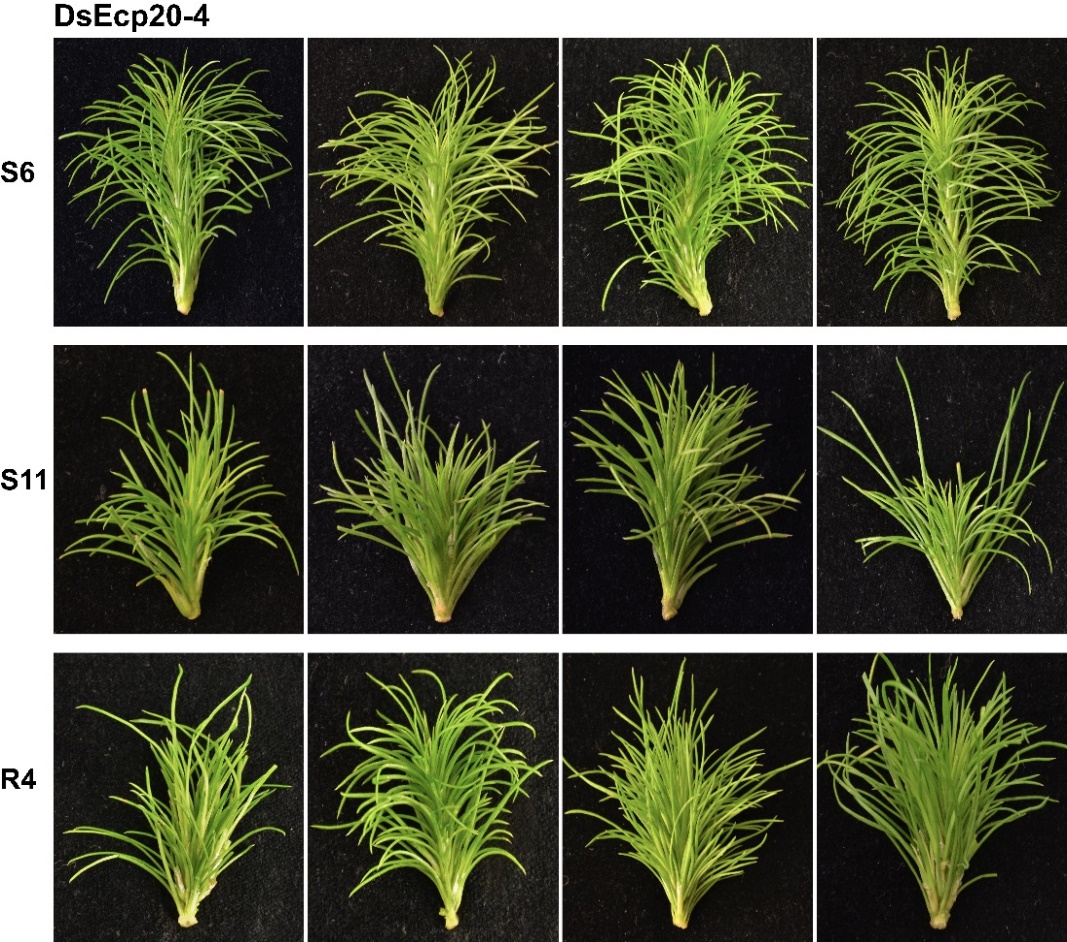

**Supplementary Figure S10.** Replicates of *Pinus radiata* shoot tissue infiltrated with DsEcp20-3 and DsEcp20-4 proteins. DsEcp20-3 (A) and DsEcp20-4 (B) were produced by heterologous expression in *Pichia pastoris*. Photos were taken 7 days after infiltration.
